# Non-invasive neuromodulation of sub-regions of the human insula differentially affect pain processing and heart-rate variability

**DOI:** 10.1101/2023.05.05.539593

**Authors:** Wynn Legon, Andrew Strohman, Alexander In, Katelyn Stebbins, Brighton Payne

**Author notes:** Wynn Legon ( / 1 Riverside Circle, Roanoke, VA, 24016, USA).

## Abstract

The insula is a portion of the cerebral cortex folded deep within the lateral sulcus covered by the overlying opercula of the inferior frontal lobe and superior portion of the temporal lobe. The insula has been parsed into sub-regions based upon cytoarchitectonics and structural and functional connectivity with multiple lines of evidence supporting specific roles for each of these sub-regions in pain processing and interoception. In the past, causal interrogation of the insula was only possible in patients with surgically implanted electrodes. Here, we leverage the high spatial resolution combined with the deep penetration depth of low-intensity focused ultrasound (LIFU) to non-surgically modulate either the anterior insula (AI) or posterior insula (PI) in humans for effect on subjective pain ratings, electroencephalographic (EEG) contact head evoked potentials (CHEPs) and time-frequency power as well as autonomic measures including heart-rate variability (HRV) and electrodermal response (EDR). N = 23 healthy volunteers received brief noxious heat pain stimuli to the dorsum of their right hand during continuous heart-rate, EDR and EEG recording. LIFU was delivered to either the AI (anterior short gyrus), PI (posterior longus gyrus) or under an inert sham condition time-locked to the heat stimulus. Results demonstrate that single-element 500 kHz LIFU is capable of individually targeting specific gyri of the insula. LIFU to both AI and PI similarly reduced perceived pain ratings but had differential effects on EEG activity. LIFU to PI affected earlier EEG amplitudes around 300 milliseconds whereas LIFU to AI affected EEG amplitudes around 500 milliseconds. In addition, only LIFU to the AI affected HRV as indexed by an increase in standard deviation of N-N intervals (SDNN) and mean HRV low frequency power. There was no effect of LIFU to either AI or PI on EDR or blood pressure. Taken together, LIFU looks to be an effective method to individually target sub-regions of the insula in humans for site-specific effects on brain biomarkers of pain processing and autonomic reactivity that translates to reduced perceived pain to a transient heat stimulus. These data have implications for the treatment of chronic pain and several neuropsychological diseases like anxiety, depression and addiction that all demonstrate abnormal activity in the insula concomitant with dysregulated autonomic function.

## INTRODUCTION

Current non-invasive neuromodulatory approaches like transcranial magnetic stimulation (TMS) and transcranial electric stimulation (TES) induce transient plastic changes in human cortex that have significantly contributed to our understanding of the human brain. However, these technologies have critical limitations including poor spatial resolution, a depth-focality tradeoff and significant attenuation at depth leading to an inability to stimulate deep neural structures with any specificity[1–4]. Low-intensity focused ultrasound (LIFU) is an emerging neuromodulatory approach that uses mechanical energy to non-destructively and reversibly modulate neuronal activity with high spatial resolution and adjustable depth of focus [5–11]. LIFU has been used safely and effectively for cortical and sub-cortical neuromodulation in mouse [12,13], rat [14,15], rabbit[16], sheep[17], pig[18], primate[19–23] and human [5,6,24,25,25–29]. LIFU is particularly advantageous for non-invasive targeting of deep neural structures in humans as it provides for millimeter lateral resolution even for deep targeting[6]. One highly promising target for pain modulation is the insula [30–34]. The insula is a portion of the cerebral cortex folded deep within the lateral sulcus covered by the overlying opercula of the inferior frontal lobe as well as the superior portion of the temporal lobe. Multiple lines of evidence demonstrate the insula as a critical brain area for nociception and the pain experience [31,33,35–41]. The insula can be broadly parsed into anterior (AI) and posterior (PI) portions based upon cytoarchitectonics, and structural and functional connectivity[42–46] and are proposed to code different aspects of the pain process[33,35]. The PI receives the majority of direct di-synaptic spino-thalamic projections[33,47] and encodes sensory aspects of nociceptive input including intensity, somatotopy and modality[30,48–51]. The AI has strong reciprocal connections with PI[35] and also to limbic regions[52–54] and is a part of the salience network[55,56]. It is postulated that PI first receives nociceptive input and relays it to AI[35] that integrates expectation, awareness and emotion to assign significance and form the overall subjective percept of pain[54].

In addition to its role in pain processing, the insula is also part of a central autonomic network[57,58] and is considered the primary interoceptive cortex[59]. The posterior insula receives afferents including vagal, glossopharyngeal and spinothalamic information from Lamina I of the spinal cord conveying information on the physiological condition of the body[59]. It is postulated that this information is used to respond to stressors (allostasis) to maintain body homeostasis[59] and that dysfunction of this interoceptive monitoring system could underlie several disease states[60,61] to which the insula looks to be a common factor[62–64]. Part of the insular role in interoception involves monitoring and control of cardiorespiratory function[65,66]. Indeed, direct electrical stimulation of either the anterior or posterior insula can affect cardiac function[67] including metrics of heart-rate variability indexing sympathetic and parasympathetic tone[68].

Based upon findings that the evoked potential from a heat stimulus is differentially recorded in AI vs. PI[35,54,69] and that sympathetic and parasympathetic control of the heart has also been found to be differentially controlled by AI vs. PI[68] we sought to investigate how and if non-invasive LIFU to either the AI or PI would differentially affect the amplitude of the contact heat evoked potential and/or heart rate variability. Given the established role of the insula in several diseases including chronic pain[59,70], addiction[61,71] and neuropsychological disease[62–64,72] and the renewed interest in the role of interoception in brain disease[60,61,73,74], the ability to modulate the insula non-invasively with high spatial precision could have broad spectrum application to the treatment of several neuropsychological diseases.

## MATERIALS & METHODS

### Participants

The Virginia Tech Institutional Review Board approved all experimental procedures. A total of N = 23 healthy volunteers (27 years ± 5.5 years age; range (19-45); M/F 7/16) provided written informed consent to participate in all aspects of the study and were financially compensated for their participation. Exclusion criteria included contraindications to other forms of non-invasive neuromodulation as outlined by Rossi et al.[75] for transcranial magnetic stimulation. Addition exclusion criteria included: contraindications to MRI; contraindications to CT including pregnancy; an active medical disorder or treatment with potential CNS effects (e.g. Alzheimer’s); a history of neurologic disorder (e.g. Parkinson’s, Epilepsy, or Essential Tremor); a history of head injury resulting in loss of consciousness for >10 minutes and a history of alcohol or drug dependence.

### Study design

This was a double-blind sham-controlled cross-over design collected over 4 sessions on 4 separate days separated by at least 2 days. Day 1 consisted of anatomical magnetic resonance imaging (MRI) and computed tomography (CT) scanning as well as baseline questionnaires. Sessions 2 – 4 were the randomized counter-balanced interventions of either LIFU to anterior insula (AI), posterior insula (PI) or Sham.

### Questionnaires

During the Day 1 anatomical imaging visit, all participants completed the following questionnaires: Beck Depression Inventory II (BDI II)[76], Beck Anxiety Inventory (BAI)[77], State-Trait Anxiety Inventory (STAI)[78], Pain Catastrophizing Scale (PCS)[79], Medical Outcomes Survey Short Form-8 (SF-8), Sleep Scale from the Medical Outcomes Study, Patient Health Questionnaire - 2 (PHQ-2)[80], Generalized Anxiety Disorder 2-item (GAD-2), Tobacco, Alcohol, Prescription medications, and other Substances Tool (TAPS), Perceived Stress Scale (PSS)[81], and the International Physical Activity Questionnaire (IPAQ). On formal testing days (Sessions 2 – 4) participants completed questionnaires both before and after the intervention. Pre session questionnaires included the STAI (state component only), the Daily Questionnaire and report of symptoms (ROS). Post session surveys included the Daily Questionnaire, Experience Questionnaire, and report of symptoms. The Daily Questionnaire queried participants on the use of substances with quantities (caffeine, nicotine, alcohol, recreational drugs) or prescription medications not previously reported that day. It also queried participants on total minutes of physical activity with intensity level (none, mild, moderate, somewhat heavy, vigorous) and three 5-point Likert scales ranging from “Disagree” to “Agree”: “I am anxious today,” “I am stressed today,” and “I slept well last night.” The Experience Questionnaire asked participants: “I could hear LIFU”, “I could feel LIFU”, “I believe I received LIFU”. The Report of Symptoms questionnaire queried symptoms (headache, sleepiness etc.) and severity (absent, mild, moderate, severe) both before and after LIFU as previously used in Legon et al. (2020)[82].

### MRI and CT Imaging

MRI data were acquired on a Siemens 3T Prisma scanner (Siemens Medical Solutions, Erlangen, Germany) at the Fralin Biomedical Research Institute’s Human Neuroimaging Laboratory. Anatomical scans were acquired using a T1-weighted MPRAGE sequence with a TR = 1400 ms, TI = 600 ms, TE = 2.66 ms, flip angle = 12°, voxel size = 0.5×0.5×1.0 mm, FoV read = 245 mm, FoV phase of 87.5%, 192 slices, ascending acquisition. Computerized Tomography (CT) scans were collected with a Kernel = Hr60 in the bone window, FoV = 250 mm, kilovolts (kV) = 120, rotation time = 1 second, delay = 2 seconds, pitch = 0.55, caudocranial image acquisition order, 1.0 mm image increments for a total of 121 images and scan time of 13.14 seconds.

### Contact Heat Evoked Potentials (CHEP)

Participants were seated in a comfortable chair with support for both arms. 40 CHEP stimuli (300 msec duration; trapezoidal stimulus with 50 msec rise/fall time at 300°C/s and a 200 msec plateau time; 32°C starting temperature; **Figure 1A**) were delivered to the dorsum of the right hand using a contact 3×3.2×2.4 mm peltier device (T03, QST.lab, Strasbourg, FR). Stimuli were delivered at a random inter-stimulus interval (ISI) ranging from 12 – 20 seconds to 6 different sites on the back of hand spaced in a 2 x 3 cm grid to help mitigate skin irritation and/or habituation. Prior to formal testing participants underwent heat pain thresholding using the same stimulus parameters as above with method of limits to determine the stimulus temperature that reliably elicited a perceived pain response of 5/9 on a 0 – 9 visual analog scale where 0 was no pain, 1-3 was mild pain; 4 – 6 was moderate pain and 7-9 severe pain. This pain scale was present on a screen in front of patient throughout testing as a reminder of the numerical scale, however there were no explicit instructions of where the participant should fixate. Participants were requested to keep their eyes open throughout testing. Head movement was limited due to the attachment of the ultrasound transducer. Participants were required to rate the perceived pain on the 0 – 9 pain scale using a numerical keypad with their left hand after feeling the CHEP stimulus. No time limits or other explicit instructions were given. Total testing time was roughly 10 – 12 minutes.

**Figure 1.**
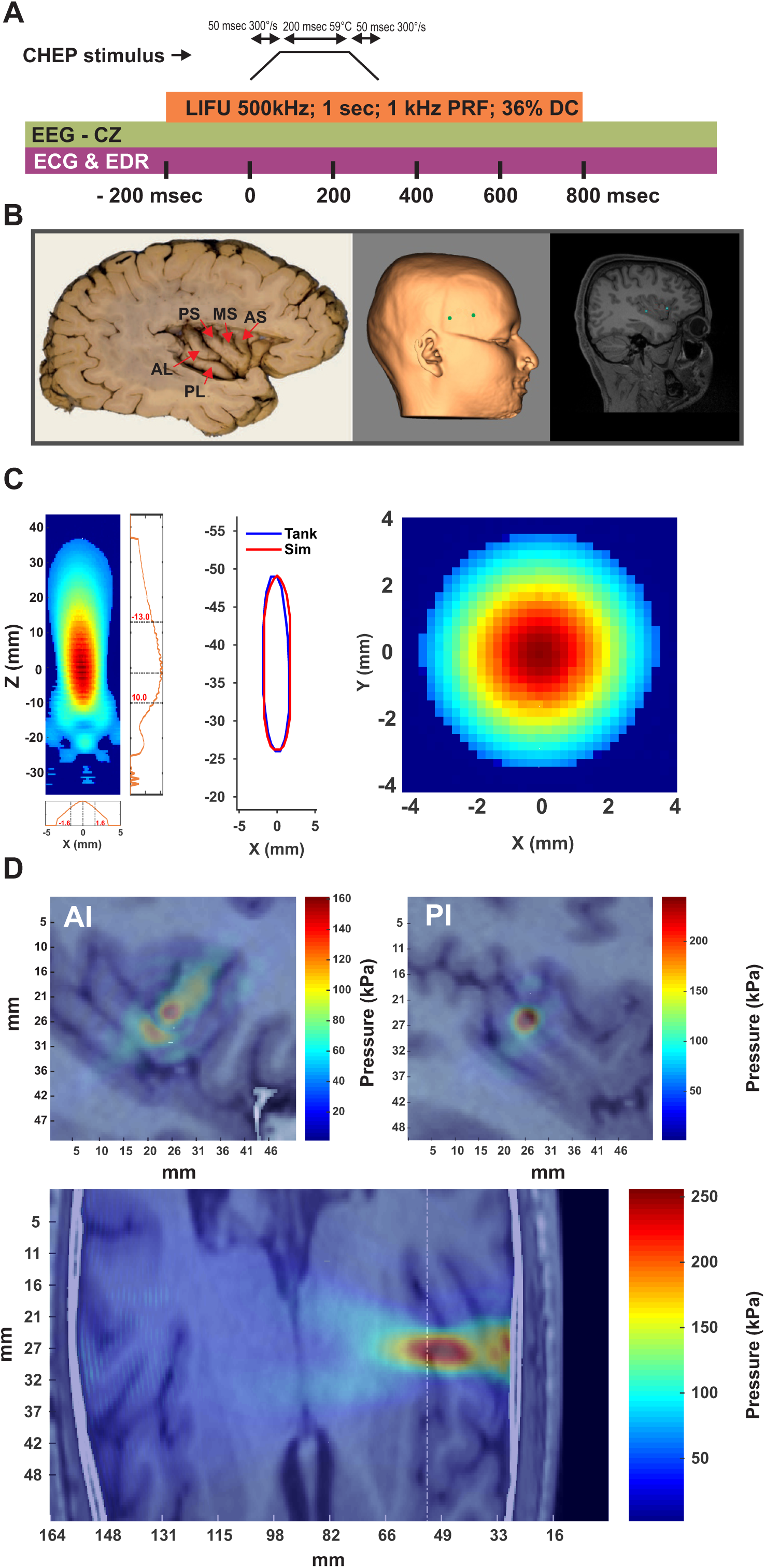
Experimental design and LIFU targeting. **A.** Pictorial representation of the timing of the contact heat evoked potential (CHEP) stimulus relative to the timing of low-intensity focused ultrasound (LIFU) and electroencephalography (EEG), electrocardiography (ECG) and electrodermal reactivity (EDR). One second of LIFU was delivered 200 msec prior to the delivery of a 300 millisecond trapezoidal CHEP stimulus to the dorsum of the right hand. **B** (left). Photograph of the individual gyri of the insula (adapted from Craig). Posterior insula is comprised of the anterior longus gyrus (AL) and posterior longus gyrus (PL). The anterior insula is comprised of the posterior short gyrus (PS), middle short gyrus (MS) and anterior short gyrus (AS). (Middle). LIFU was targeted to the dorsal aspect of the AS and AL. Targets are shown in green on a scalp rendering of one participants. (Right) Targets shown on same participants’ sagittal MR image. **C.** (Left) Pseudocolor axial (XZ) empirically measured LIFU beam from acoustic test tank. Numbers in red represent limits of the full-width half-maximum pressure in millimeters from the focal spot. (Middle) Overlay of the FWHM of the empirical measurements (Tank) with the modelled beam used for acoustic modelling (Sim). (Right) Pseudocolor lateral (XY) empirically measured LIFU beam. The FWHM lateral resolution is ~ 3.4 mm. **D.** (Top) Sagittal views of acoustic models showing targeting of anterior insula (AI) and posterior insula (PI) in one representative subject. (Bottom) Transverse view showing beam targeting anterior insula. White line represents plane from which AI top image was taken.

### Electroencephalogram (EEG)

Data were acquired using a DC amplifier (GES 400, Magstim EGI, Eugene, OR, USA) and two 10-mm silver-silver chloride cup electrodes placed at the vertex (Cz) and the central frontal location (Fz) referenced to the bilateral mastoid. Prior to electrode placement, the scalp was first prepared with a mild abrasive gel (Nuprep; Weaver and Company, Aurora, CO) and then rubbing alcohol. Cup electrodes were filled with a conductive paste (Ten20 Conductive; Weaver and Company, Aurora, CO) and held in place with medical tape. Electrode impedances were verified (<50 kΩ) before recording. Data were continuously sampled at 1 kHz using a 64-channel EEG recording system (GES 400, Magstim EGI, Eugene, OR, USA) and Net Station™ 5.4 EEG software and stored on a PC for offline data analysis.

### Physiological recordings

#### Electroencephalogram (ECG)

Heart-rate was collected using dual lead ECG with one electrode attached to the anterior surface of each forearm immediately distal to the antecubital fossa. Data was continuous sampled at 1 kHz using the Physio16 input box and EEG recording system and Net Station™ 5.4 EEG software (GES 400, Magstim EGI, Eugene, OR, USA). *Respiration*. Respiration data was collected using a piezoelectric respiration sensor (gTec™ gSensor Respiration Effort) over the participants clothing placed over the lower ribs outside of clothing. Respiration data was continuously sampled at 1 kHz using the same hardware and software setup as ECG above. *Electrodermal Response (EDR).* EDR was collected using the Consensys™ GSR Development Kit (Shimmer, Cambridge, MA, USA). It was placed on the right wrist with a photoplethysmogram (PPG) sensor on the distal 4^th^ digit and two EDR electrodes on the distal 2^nd^ and 3^rd^ digits. EDR data was continuously collected and sampled at 128 kHz using the Consensys™ software and stored on a PC for offline data analysis. *Blood Pressure (BP).* Systolic and diastolic blood pressure was collected using the QardioArm™ wireless portable blood pressure monitor placed on the upper left arm. BP measurements were taken twice; once at the beginning of each session prior to the intervention and again at the end of each session (see **Figure 1A** for schematic of timing of stimulation and recording).

### LIFU Transducer

We used a Sonic Concepts H-281 single-element 500 kHz transducer with an active diameter of 45.0 mm and a geometric focus of 45.0 mm. The focal depth from the exit plane was 38.0 mm. The transducer also had a solid water coupling over the radiating surface to the exit plane.

### LIFU waveform

LIFU waveforms were generated using a two-channel, 2-MHz function generator (BK 4078B Precision Instruments). Channel 1 was used to gate channel 2 that was a 500 kHz sine wave. Channel 1 was a tapered 5Vp-p square wave burst of 1 kHz (N = 1000) with a pulse width of 360 msec. This resulted in a 1.0 sec duration waveform with a duty cycle of 36%. The output of channel 2 was sent through a 100-W linear RF amplifier (E&I 2100L; Electronics & Innovation) before being sent to the LIFU transducer. The waveform was time-locked to occur 200 msec prior to the CHEP stimulus. The peak negative pressure of the waveform outside the head was 400 kPa or 3.5 W/cm^2^ spatial peak pulse average intensity (Isppa) for all participants.

### LIFU targeting

The transducer was coupled to the head using conventional ultrasound gel and our custom mineral oil/polymer coupling pucks[83]. These pucks have negligible attenuation at 500 kHz[83] and can be made with varying stand-off heights that allow for precise axial (depth) targeting based on individual insular target depths. Each participants’ AI and PI target was identified with the aid of an insular atlas[42] and depth measured from the scalp. An appropriate coupling puck was made so that the focal spot of the transducer (38 mm) exactly overlaid on the insular target (see **Table 1** & Results for insula target depths and MNI co-ordinates). Placement of the transducer on the scalp was aided using a neuronavigation system (BrainSight, Rogue Research, Montreal, QUE, CAN). Prior to formal testing, the AI (dorsal aspect of the anterior short gyrus) and PI (dorsal aspect of the anterior longus gyrus) were identified on individual subject anatomical MRI scans. These co-ordinates were entered into the neuronavigation system and used for online tracking of transducer placement throughout the testing session. LIFU or Sham was only delivered if placement error was < 3 mm (see Results for targeting error and see **Figure 1B** for example of neuronavigated targeting).

### Acoustic masking

In some cases, a single-element transducer can produce an audible auditory artifact likely as the result of the pulse repetition frequency. In order to remove this as a potential confound[84], strict acoustic masking was performed. Acoustic masking was delivered through disposable earbuds that were plugged into a Kindle tablet. Participants were instructed to select a combination of sounds from a white noise app that were then randomly mixed creating a multitone. Participants were told to set the volume to something that was comfortable yet removed ambient sounds. The intensity range was on average 70 – 75 dB. Auditory masking was confirmed by speaking to the participant out of their visual field. Auditory masking noise was played continuously throughout the testing session. 30 minutes after formal testing, participants were queried on auditory masking. Questions included “I could hear the LIFU stimulation”, “I could feel the LIFU stimulation”, and “I believe I experienced LIFU stimulation.” Participants were requested to respond to each question using a 7-point Likert scale (0 – 6) with points corresponding to Strongly Disagree / Disagree/ Somewhat Disagree / Neutral / Somewhat Agree / Agree / Strongly Agree.

#### Skull density ratio (SDR) and thickness

Skull density ratio (SDR) is the ratio of cortical to trabecular bone (as measured in Hounsfield units from CT) and has been demonstrated to be predictive of the success of high-intensity focused ultrasound ablative surgery[85,86]. To calculate SDR and skull thickness, we used individual participant MR and CT scans. MR scans were used to determine a spot on the scalp equidistant between the AI and PI scalp targets. We then used individuals participants’ co-registered CT scans to get the Hounsfield units from the skull from an area +10 mm to −10 mm from the target spot in the dorsal/ventral and anterior/posterior axes (0.4588 mm resolution = 79 total samples). Cortical bone values were taken as the average of the first and second maxima and trabecular bone values were taken as the average of the points between these two values. Skull thickness was calculated as the absolute distance between points where the Hounsfield units were > 100. All SDR and skull thickness data was analyzed using custom script written in Matlab® v.9.5 (R2018b) (The MathWorks Inc., Natick, MA, USA).

### Analysis

#### Questionnaires

Data from the auditory masking questionnaires (0 – 6 pts) were each subjected to non-parametric kruskalwallis tests (AI, PI and Sham). Data from the BDI, STAI and PCS are reported as mean ± SD.

#### EEG

Only data from the CZ electrode were quantified as this electrode displayed the clearest CHEP waveform for all participants. EEG data were preprocessed using custom scripts written in Matlab® v9.5.0 (R2018b) (The MathWorks, Inc., Natick, MA). Data were band-pass filtered (2–100 Hz) using a third-order Butterworth filter and the filtfilt function in Matlab®, data were then epoched around the CHEP stimulus (−2000 to 2000 msec) and baseline corrected (−1500 to −500 msec). Data were manually inspected for artifact (eye blink, muscle activity) and contaminated epochs removed. The maximum number of rejected trials for a single participant was 1. Two forms of CHEP waveform analysis were employed: time series analysis and peak-to-peak analysis. Time series analysis provides for investigation of not just the CHEP peaks but for any differences across time between the conditions of interest. For time series analysis, traces (0 – 1000 msec) from channel CZ were analyzed using nonparametric permutation statistics (p<.05; 5,000 randomizations) which appropriately control for multiple comparisons problems encountered in analyses of complex EEG data sets[87], where statistical P values represent the proportion of 5,000 random partitions resulting in a F-statistic larger than the F value calculated by a conventional one-way repeated measures ANOVA (AI, PI, Sham) on the data. In addition, a temporal cluster threshold of at least 10 consecutive time-points (10 msec) was required to satisfy significance. For statistically significant data points, main effects were investigated using permutation statistics using t-statistics for each of the tests: AI vs. PI, AI vs. Sham and PI vs. Sham. Significance was set at p < 0.05. For peak-to-peak analysis, waveform peak amplitude and latency were manually identified and quantified. Peaks of interest included the first large negative deflection around 330 msec (N1) and the first large positive deflection around 500 msec (P1)[88,89]. A distinct inflection of the waveform was necessary for inclusion in statistical analyses. All participants displayed clear N1/P1 peaks above the noise floor. Statistical analyses was conducted on N1/P1 peak-to-peak amplitudes as well as individual N1 and P1 amplitudes using separate one-way repeated measures analysis of variance (ANOVA) with main factor LIFU (AI, PI, Sham). Statistical significance was set at p < 0.05. Significant main effects were post-hoc tested using Tukey-Kramer tests (p < 0.05). All statistical analysis was performed in Matlab® using built-in functions and custom scripts.

#### EEG time-frequency spectra

Average CHEP epochs from −2000 msec to + 2000 msec centered around the CHEP stimulus for each condition for each individual were convolved with a Morlet wavelet using 30 frequencies log-spaced between 2 – 100 Hz. To assess non-temporally specific mean power changes across conditions, data from the time window 200 to 800 msec were grouped into canonical EEG frequencies including delta (2-4 Hz), theta (4-8 Hz), alpha (8-13 Hz), beta (14 – 30 Hz), low gamma (30 – 60 Hz) and high gamma (60 – 100 Hz) and normalized for each condition by dividing by that conditions’ baseline power (−1500 msec to −200 msec). The 200 – 800 msec time window was chosen as this looked to be the time window that best captured meaningful power fluctuations as a result of the noxious stimulus. The mean power for each normalized power band was compared across conditions (AI, PI and Sham) using separate one-way repeated measures ANOVA. In a separate analysis, to better understand the timing of these changes we also performed non-parametric permutation statistics (5000 permutations, p < 0.05, cluster threshold of 10 consecutive time points (10 msec)) across for each frequency (30 log-spaced frequencies 2 – 100 Hz) across the time window −100 to 1000 msec where 0 is the timing of the noxious stimulus.

#### Heart rate

Heart rate data was first filtered from 10 – 30 Hz using a 3^rd^ order Butterworth filter and Matlab® function filtfilt. Manual inspection of the data stream was then performed and any artifact epochs removed. R peaks were extracted using the findpeaks function in Matlab® and manually confirmed. The entire data stream (~ 25 minutes: whole testing session which includes resting periods) was epoched into time windows prior to LIFU (~ 5 minutes resting), during LIFU (duration : ~ 12 – 15 minutes) and after LIFU (~ 5 minutes resting). Only data from the epoch during the LIFU/CHEPs is analyzed and presented here.

#### Electrodermal Response (EDR)

EDR data was bandpass filtered (0.1 – 25 Hz) using a 3^rd^ order butterworth filter and filtfilt Matlab function. The mean of the entire EDR time series was removed and then data was epoched around the onset of the CHEP stimulus (−5 to 10 seconds) and averaged for the 40 CHEPs trials for each condition for each individual. EDR response was quantified as the peak-to-peak (absolute of min-max) of the waveform in the interval after the CHEP onset (0 – 10 seconds). These data were analyzed using a one-way repeated measures ANOVA with factors (AI, PI, Sham).

#### Acoustic Modelling

Computational models were developed using individual subject MR and CT images to evaluate the wave propagation of LIFU across the skull and the resultant intracranial acoustic pressure maps. Simulations were performed using the k-Wave MATLAB toolbox[90], which uses a pseudospectral time domain method to solve discretized wave equations on a spatial grid. CT images were used to construct the acoustic model of the skull, while MR images were used to target LIFU at either the AI or PI target, based on individual brain anatomy. Details of the modelling parameters can be found in Legon et al. (2018)[6]. CT and MR images were first co-registered and then resampled for acoustic simulations at a finer resolution and the acoustic parameters for simulation calculated from the CT images. The skull was extracted manually using a threshold intensity value and the intracranial space was assumed to be homogenous as ultrasound reflections between soft tissues are small[91]. Acoustic parameters were calculated from CT data assuming a linear relationship between skull porosity and the acoustic parameters[92,93]. The computational model of the ultrasound transducer used in simulations was constructed to recreate empirical acoustic pressure maps of focused ultrasound transmitted in the acoustic test tank similar to previous work[91].

## RESULTS

### Ultrasound beam characteristics

The ultrasound beam as measured in free water had a lateral full-width at half maximum (FWHM) resolution in the X plane of 3.3 mm and in the Y plane of 3.4 mm at the Z maximum. The axial FWHM was 23 mm ranging from −10 mm to + 13 mm from the point of maximum pressure (38 mm from exit plane) conferring and effective axial FWHM of 28 – 51 mm (see **Figure 1C**). The constructed model waveform used for all acoustic simulations was in good agreement with these empirical measurements validating its use in the models (**Figure 1C**).

### Insula targets

The MNI co-ordinates for AI and PI for each participants are in **Table 1**. The average depth of the AI target for males and females was: 35 ± 3.7 mm and 34.2 ± 2.5 mm respectively. The depth of the PI target for males and females was: 40.3 ± 3.4 mm and 39.4 ± 2.1 mm (see **Table 1**). These values overlap with the empirical and modelled FWHM of the waveforms that encompassed an axial depth FWHM resolution from 28 mm to 51 mm (see **Figure 1D**). The average distance between targets (anterior to posterior) for the group (N = 23) was 25.9 mm ± 3.3 mm. For males and females it was: 28.0 mm ± 3.9 mm and 24.9 mm ± 2.6 mm respectively (see **Figure 1D** for acoustic models of AI and PI).

### Temperature of CHEP stimulus

The average temperature of the Peltier device to elicit a 5/9 perceived pain rating for AI, PI and Sham conditions was: 59.5 ± 1.2°C, 59.6 ± 1.2°C and 59.3 ± 1.5°C. For males it was 59.9 ± 0.4, 59.9 ± 0.4 and 59.7 ± 0.8 respectively. For females it was: 59.3 ± 1.4, 59.5 ± 1.4 and 59.1 ± 1.7 respectively. We conducted a two-way mixed ANOVA with factors SEX (M, F) and LIFU (AI, PI, Sham). There was no main effect of SEX F(1,68) = 2.02, p = 0.16 or LIFU F(2,68) = 0.19, p = 0.83 and no interaction F(2,68) = 0.04, p = 0.96.

### Perceived Pain

The mean of 40 individual responses to the CHEP stimuli were averaged for each subject for each condition (AI, PI, Sham). Mean ± SE for AI, PI and Sham were: 3.03 ± 1.42; 2.77 ± 1.28 and 3.39 ± 1.09. This data was subjected to a one-way repeated measures ANOVA. This revealed a significant main effect F(2,44) = 4.29, p = 0.019. Post-hoc Tukey test (p < 0.05) revealed the main effect was driven by a significant difference between posterior insula (PI) and Sham stimulation. There was no significant statistical difference between AI and Sham or AI and PI (see **Figure 2A**).

**Figure 2.**
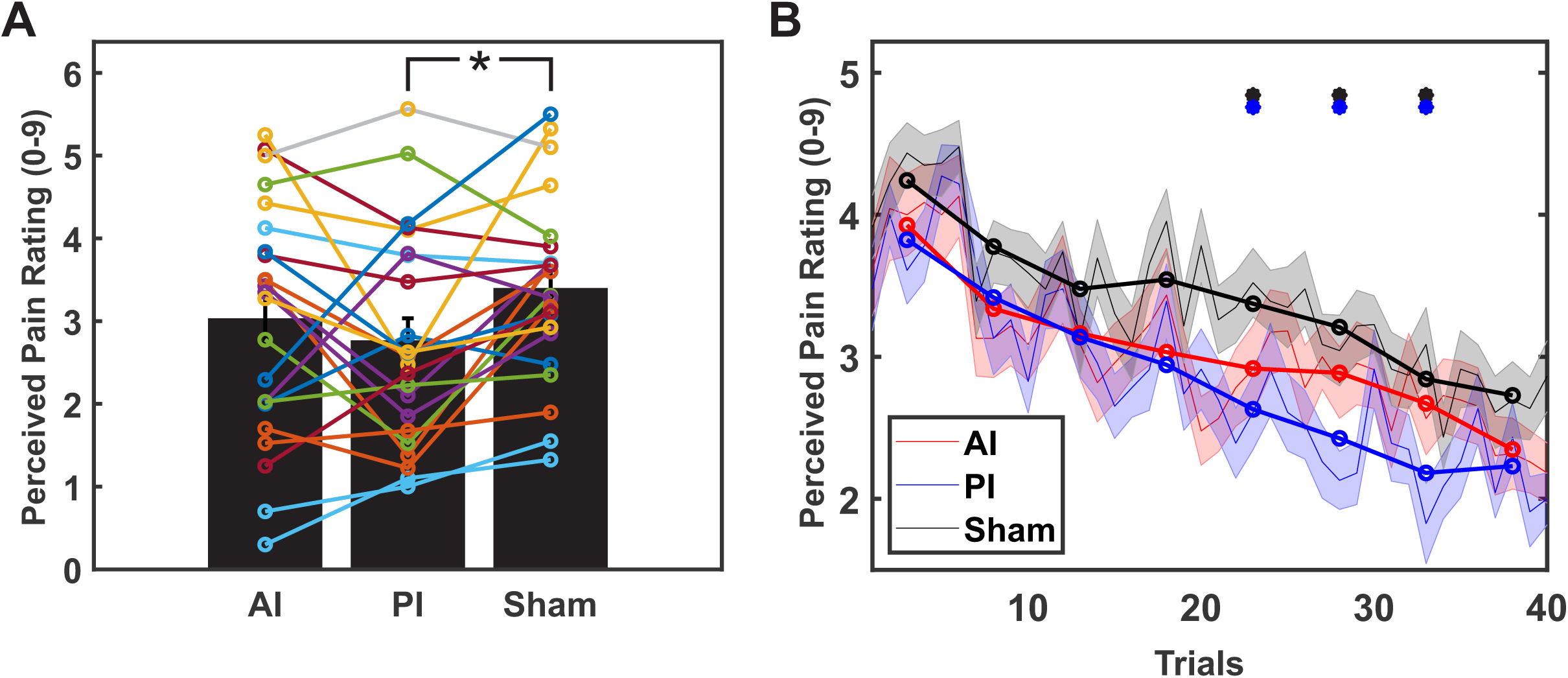
Effects of LIFU on behavior. **A.** Grand average (N = 23) perceived pain ratings to a brief contact heat stimulus to the dorsum of the hand during LIFU to either anterior insula (AI), posterior insula (PI) or Sham stimulation. Bars are mean ± standard error of the mean (SEM). Individual subject data is overlaid. * denotes p < 0.05. **B.** Grand average (N = 23) perceived pain ratings over the 40 trials (~ 10 minutes). Thin shaded lines represent mean ± SEM of individual trials. Thicker lines with circles represent the average of every 5 trials. LIFU to anterior insula (AI), posterior insula (PI) and Sham are shown in red, blue and black respectively. Dots represent statistically significant differences (p< 0.05 permutation statistics) between Sham (black) and PI (blue).

### Perceived Pain Temporal effects

To investigate the effect of repeated LIFU stimulations on perceived pain perception we binned the 40 trials into 8 bins of the mean of 5 trials (1-5,6-10,11-15,16-20,21-25,26-30,31-35,36-40) for each participant for each condition (AI, PI, and Sham). For this analysis we ran non-parametric permutation statistics (p < 0.05, 5000 randomizations). Tests which resulted in a p < 0.05 were further investigated using post-hoc Tukey tests to investigate what differences were driving the main effect. Permutation statistics revealed bins 5, 6 and 7 demonstrated significant effects (p < 0.05). Post-hoc Tukey tests revealed the main effect was driven by differences between PI and Sham for all significant bins. Of note, the first bin did not reveal a statistically significant difference (p = 0.27) suggesting that the initial starting point of perceived pain rating was not different between groups. The mean ± SD for the first bin for AI, PI and Sham was: 3.93 ± 1.82; 3.82 ± 1.3 and 4.24 ± 1.23 respectively (see **Figure 2B**).

### CHEP peaks

#### N1/P1 peak-to-peak results

Mean ± SD N1/P1 amplitudes for AI, PI and sham were: 23.35 µV ± 11.58 µV; 22.90 µV ± 12.35 µV and 27.79 µV ± 10.78 µV. The one-way repeated measures ANOVA revealed a main effect F(2,44) = 8.19, p = 0.009. Post-hoc Tukey tests revealed significant difference between Sham and AI (p<0.05) and Sham and PI (p<0.05). There was no significant difference between AI and PI (**Figure 3A**). Peak-to-peak analysis does not inform on if either the N1 or P1 was more or less affected by the intervention. As such, we also looked at the N1 and P1 peak amplitudes separately.

**Figure 3.**
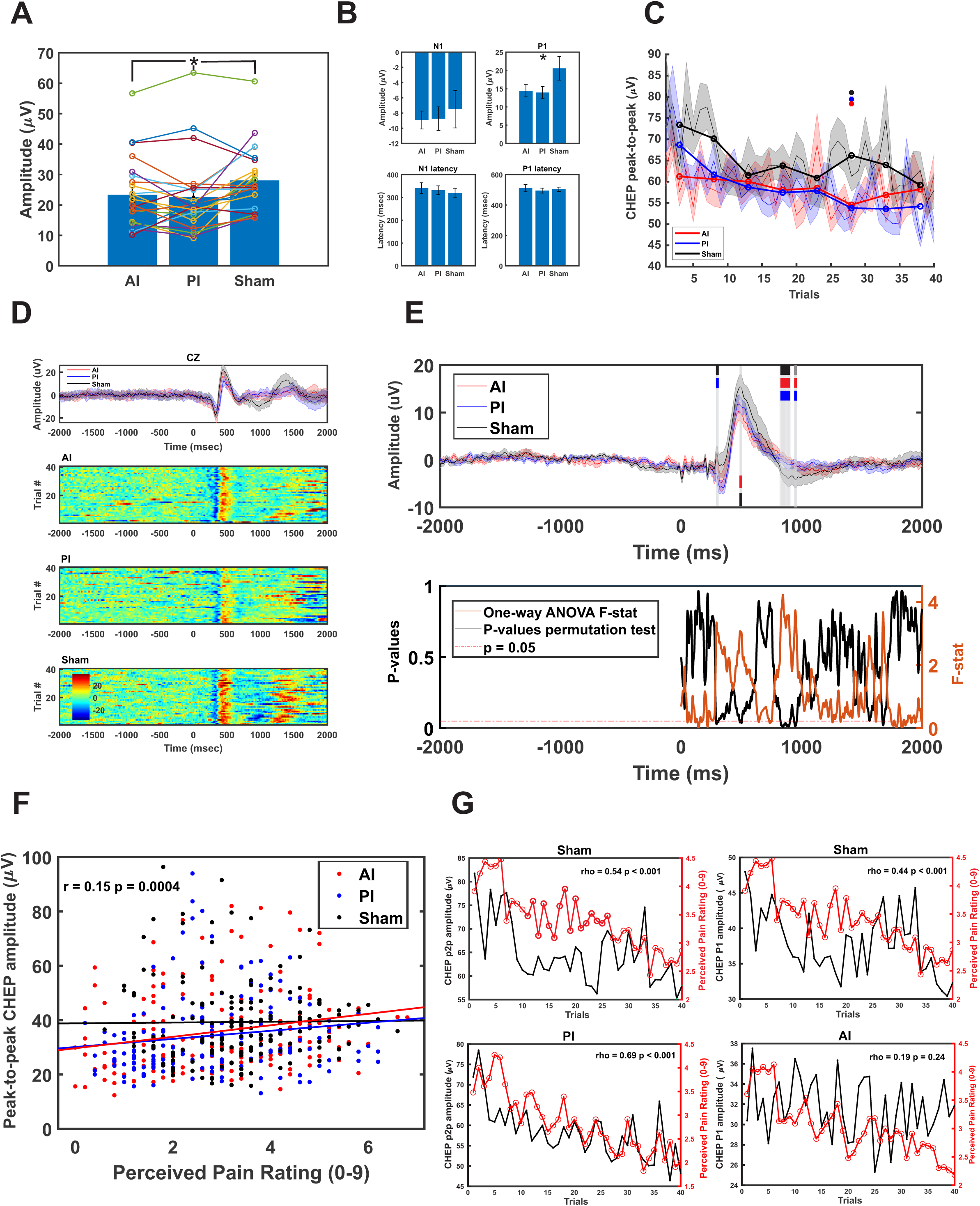
Effect of LIFU on the contact heat evoked potential (CHEP). **A.** Grand average (N = 23) peak-to-peak N1/P1 CHEP amplitudes for LIFU to anterior insula (AI), posterior insula (PI) and Sham stimulation. Bars are mean ± SEM. Individual subject data is overlaid on top. * denotes significant difference (p< 0.05) for AI and PI compared to Sham. **B.** Grand average (N = 23) individual peak (N1 and P1) amplitudes and latencies. Bars are mean ± SEM. * denotes significant difference (p<0.05) between Sham and both AI and PI. **C.** Grand average (N = 23) CHEP peak-to-peak amplitude over the 40 trials (~ 10 minutes). Thin shaded lines are mean ± SEM from each trial. Thicker lines with circles represent the average of every 5 trials. Only the 6^th^ bin showed statistically significant differences (p < 0.05) where both AI (red) and PI (blue) were smaller than Sham (black) represented as dots over 6^th^ bin. **D.** Date from one representative subject showing averaged CHEP trace from 40 trials (top). Pseudocolor raster plots from the 40 trials for each of AI, PI and Sham are shown below. Scales are identical for each raster plot. **E.** Grand average (N = 23) CHEP traces for AI, PI and Sham conditions. Results of the permutation statistics (5000 randomizations, p < 0.05) run across each time point (0 – 2000 msec) are shown below. The f-statistic is shown in ochre and the corresponding p-value in black. P-values < 0.05 are highlighted above in trace with gray bars. The colors in the gray bars denote which conditions were significantly different. Sham (black) and PI (blue) were statistically different around 300 msec; AI (red) and Sham (black) were statistically different around 500 msec and both AI and PI were statistically different from Sham around 950 msec. The light green marker around 1600 msec was also significant but was not further investigated. **F.** Group (N = 23) relationship between peak-to-peak CHEP amplitude and perceived pain ratings. R and p-values are for all conditions collapsed together. Data from AI, PI and Sham conditions is color-coded for display purposes and least-squares lines for each of the conditions is plotted. **G.** Group (N = 23) correlations between peak-to-peak CHEP amplitude (black) and perceived pain behavior (red) plotted over the 40 trials (~ 10 minutes). (Left) LIFU to PI (bottom) strengthens the relationship between CHEP amplitude and behavior as compared to Sham (top). (Right) LIFU to AI weakens the relationship between CHEP amplitude and behavior (bottom) as compared to Sham (top).

#### N1 results

The mean ± SD of the amplitude of the N1 for AI, PI and Sham were: −8.91 µV ± 5.68 µV; −8.85 µV ± 7.39 µV and −7.50 µV ± 11.85 µV. There was no main effect of condition on the amplitude of the N1 potential F(2,44) = 0.33, p = 0.72 (**Figure 3B**).

#### P1 results

The mean ± SD of the amplitude of the N1 for AI, PI and Sham were: 14.44 µV ± 8.06 µV; 14.06 µV ± 7.85 µV and 20.29 µV ± 15.53 µV. The one-way repeated measures ANOVA revealed a main effect F(2,44) = 3.72, p = 0.032. Post-hoc Tukey tests revealed a significant difference between PI and Sham (p <0.05) and no statistically significance differences between AI and Sham or AI and PI (**Figure 3B**).

#### Peak latencies

The peak latencies for N1 for AI, PI and Sham conditions were: 341 ms ± 113 ms; 331 ms ± 92 ms and 320 ms ± 100 ms. There were no significant differences of N1 latencies: F(2,44) = 0.36, p = 0.70. The peak latencies for P1 for AI, PI and Sham conditions were: 511 ms ± 116 ms; 497 ms ± 82 ms and 503 ms ± 70 ms. There were no significant differences of P1 latencies: F(2,44) = 0.15, p = 0.86 (**Figure 3B**).

### CHEP temporal dynamics

We investigated the temporal effects of LIFU on peak-to-peak N1/P1 amplitudes as well as just the N1 and P1. We first binned the 40 trials into 8 bins of the mean of 5 trials (1-5,6-10,11-15,16-20,21-25,26-30,31-35,36-40) for each participant for each condition (AI, PI, and Sham). We ran separate non-parametric permutation statistics (p < 0.05, 5000 randomizations) for each of N1/P1 peak-to-peak and N1 and P1. The N1/P1 peak-to-peak revealed only bin 7 to have a significant main effect (p < 0.05) that was driven by significant differences between Sham and both AI and PI (post-hoc Tukey test). No bins for N1 or P1 reached statistical significance.

Based upon these results there does not look to be temporal or cumulative effects of LIFU on the amplitude of any components of the CHEP. As a check for initial group differences, the mean of the first bin for AI, PI and Sham was: 61.25 ± 18.84; 68.69 ± 17.16 and 73.42 ± 32.43 respectively. There was no difference between groups for this bin (5000 permutations, p = 0.15) (see **Figure 3C**). Example from a single subject showing evolution of the CHEP amplitudes across trials for each condition is shown in **Figure 3D**.

### EEG time-windowed amplitude

While the CHEP peak-to-peak analysis found effects for the manually identified peaks of interest, this analysis is somewhat limited in that it does not look at differences across the entire time window. Thus, we performed an additional analysis across the time window of 0 – 2000 msec using non-parametric permutation statistics (5000 permutations, p < 0.05 with a 10 msec cluster threshold). This analysis revealed a significant main effect across condition in three separate clusters at time points 290 – 313 msec; 485 – 504 msec and 824 – 903 msec. Post-hoc testing revealed the first cluster to be driven by a significant difference between Sham and PI (p < 0.05). The second cluster was driven by a significant difference between Sham and AI only (p < 0.05). The third cluster was driven by differences between Sham and AI and Sham and PI (p < 0.05) (**Figure 3E**).

### Correlation of behavior with CHEP amplitudes

We performed separate Pearson’s correlation of the N1, P1 and N1/P1 peak-to-peak amplitudes pooled across all conditions and all trials. This analysis resulted in a significant positive correlation for peak-to-peak CHEP amplitude and perceived pain rating scores r = 0.15; p = 0.0004. Neither the N1 nor P1 alone resulted in significance (r = 0.01 and r = −0.02) (**Figure 3F**). We were also interested in the temporal evolution of the relationship between CHEP peak-to-peak, N1 and P1 amplitudes and perceived pain ratings and how LIFU to either AI or PI affected this relationship. As such, we averaged all participants (N = 23) peak-to-peak, N1 and P1 across trials (N = 40) as well as their perceived pain ratings. For the Sham condition, there was a significant positive relationship between CHEP peak-to-peak amplitude and behavior (r = 0.54, p < 0.001) (**Figure 3G**). Looking at the N1 and P1 individually, both also significantly correlated with behavior (N1: r = −0.42, p = 0.01; P1: r = 0.44, p = < 0.001). This is not surprising as the peak-to-peak is a composite variable of both the N1 and P1. Nevertheless, we further examined how and if LIFU to either AI or PI affected this relationship by statistically comparing the Sham correlations separately with the correlations during LIFU to AI and PI using Fischer’s Z transformation[94]. While LIFU to PI looked to strengthen these relationships it did not statistically significantly alter this relationship for peak-to-peak (z = −1.049, p = 0.147), N1 (z = −1.562, p = 0.059) or P1 (z = −0.691, p = 0.245). LIFU to AI looked, in general, to have the opposite effect to PI where the relationships were weakened: peak-to-peak (r = 0.36), N1 (r = −0.38) and P1 (r = 0.19) There was statistically significant difference for the P1 component (z = 1.895, p = 0.029) whereby LIFU to AI significantly disrupted or weakened the relationship between P1 amplitude and perceived pain rating (**Figure 3G**).

### EEG Time/frequency results

#### Delta (2-4 Hz)

The mean normalized power to baseline (−1500 to −500 msec) for delta from the time window 200 msec to 800 msec for AI, PI and Sham was: 1.20 ± 0.42; 1.38 ± 0.17 and 1.38 ± 0.18. The repeated measures ANOVA revealed a main effect: F(2,44) = 4.28, p = 0.02. Post-hoc Tukey tests revealed significant differences between AI and PI and AI and Sham (p<0.05) (**Figure 4A**).

**Figure 4.**
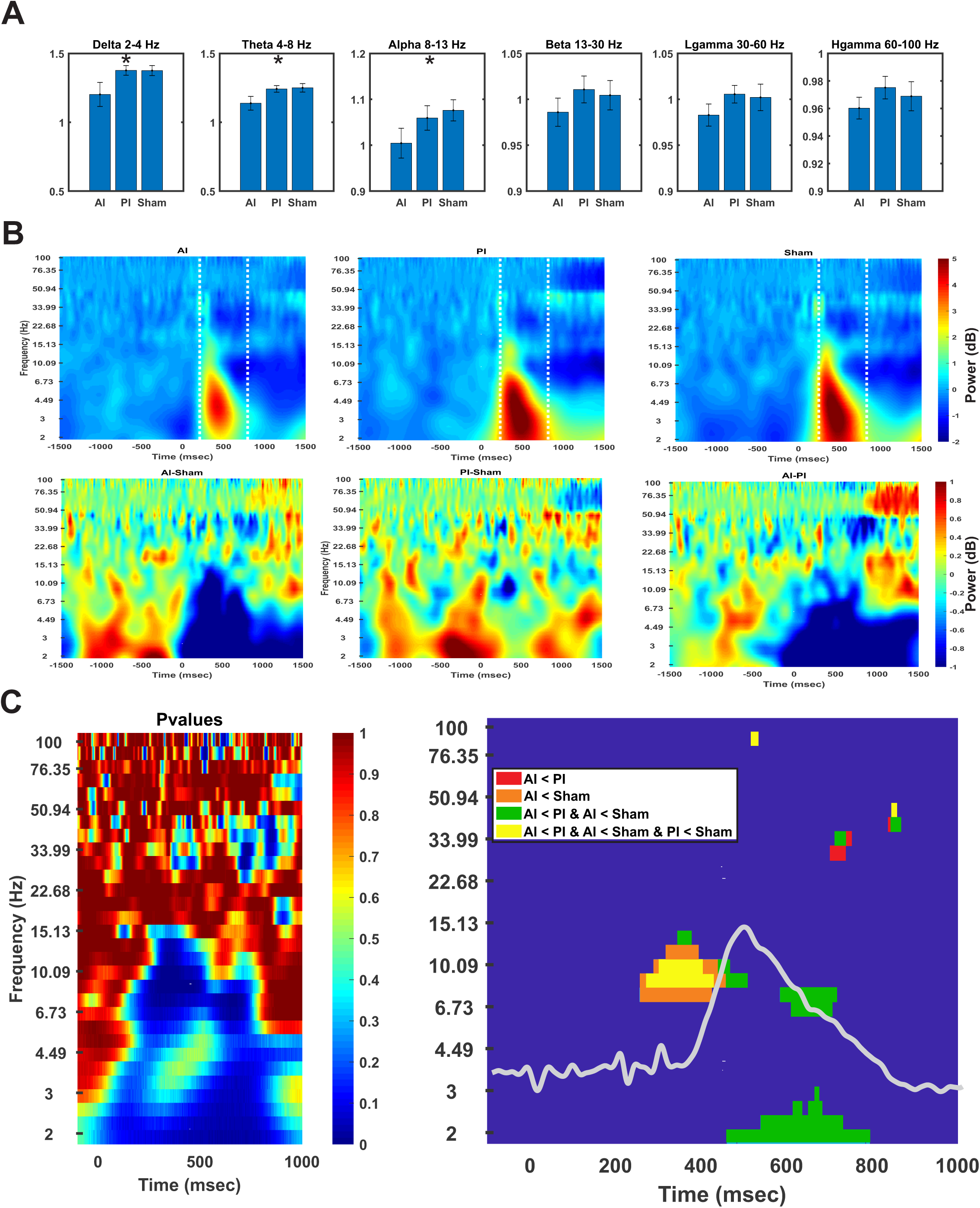
Effect of LIFU on EEG power. **A.** Group (N = 23) EEG power taken from the window 200 – 800 msec (denoted below in B by vertical white lines) relative to a baseline period. Bars are mean ± SEM. For delta and theta frequencies * denotes significant difference (p< 0.05) between both AI, PI and Sham. For alpha, * denotes significant difference (p<0.05) between AI and Sham only. **B**. (Top) Group (N = 23) mean power. (Bottom) Power difference maps taken from data above. **C.** (Left) Plot of p-values from permutation statistics (5000 randomizations). (Right) Plot showing statistically significant (p<0.05; 10 msec cluster threshold) frequency time-windows form permutation statistics. CHEP trace is overlaid in grey to help illustrate timing of frequency effects.

#### Theta (4-8 Hz)

The mean power for theta for AI, PI and Sham was: 1.14 ± 0.24; 1.24 ± 0.12 and 1.25 ± 0.14. The repeated measures ANOVA revealed a main effect: F(2,44) = 4.32, p = 0.02. Post-hoc Tukey tests revealed significant differences between AI and PI and AI and Sham (p<0.05) (**Figure 4A**).

#### Alpha (8-13 Hz)

The mean power for alpha for AI, PI and Sham was: 1.00 ± 0.16; 1.06 ± 0.13 and 1.07 ± 0.11. There was a significant main effect of condition: F(2,44) = 3.84, p = 0.03. Post-hoc Tukey tests revealed this was driven by a significant difference between Sham and AI (p < 0.05) (**Figure 4A**).

#### Beta (13-30 Hz)

The mean power for beta for AI, PI and Sham was: 0.99 ± 0.07; 1.01 ± 0.07 and 1.00 ± 0.07. There was no significant main effect of condition: F(2,44) = 1.25, p = 0.3.

#### Low-gamma (30-60 Hz)

The mean power for low-gamma for AI, PI and Sham was: 0.98 ± 0.06; 1.00 ± 0.05 and 1.00 ± 0.07. There was no significant main effect of condition: F(2,44) = 2.21, p = 0.12 (**Figure 4A**).

#### High-gamma (60 – 100 Hz)

The mean power for high gamma for AI, PI and Sham was: 0.96 ± 0.04; 0.98 ± 0.04 and 0.97 ± 0.05. There was no significant main effect of condition: F(2,44) = 1.17, p = 0.32 (**Figure 4A**).

The full-windowed (−1500 to 1500 msec) grand average (N = 23) time-frequency plots for AI, PI and Sham are shown in **Figure 7B** including difference maps for AI-Sham, PI-Sham and AI-PI.

#### Time-frequency temporal characteristics

We ran permutation statistics (5,000 randomization, p < 0.05) across each frequency band across the time window −100 to 1000 msec to better understand the temporal characteristics of frequency changes between conditions. This analysis found main effects (one-way ANOVA p < 0.05) changes in theta and alpha power clustered around the 300-400 msec window that lines up with the N1 CHEP peak. There was also a cluster of frequency change in the theta/alpha range around 600 msec. Changes in delta frequency only occurred at the timing of the P1 peak and later. Beta changes were identified later than 600 msec (see **Figure 4C**). Post-hoc analysis revealed that these main effects were largely driven by LIFU to AI. In all cases, LIFU to AI reduced power in these frequencies and time windows. **Figure 4C** color codes the post-hoc analyses.

### Autonomic data

#### Blood Pressure

The mean ± SD systolic/diastolic blood pressure for before and after LIFU for AI, PI and Sham conditions was: 119/70 ± 16/9 and 117/69 ± 14/9; 120/69 ± 13/11 and 118/70 ± 15/9; 119/69 ± 14/10 and 118/69 ± 13/10. The change scores pre/post for each of systolic and diastolic blood pressure for each condition (AI, PI, Sham) were: 1.3/0.3 ± 8.2/5.8; 1.8/-0.6 ± 6.4/4.8 and 1.4/0.0 ± 9.6/6.0. Change scores were subjected to separate one-way repeated measures ANOVAs. There were no significant differences for either systolic (p = 0.97) or diastolic (p = 0.86) blood pressure.

#### Heart rate

The mean number of R-peaks from which all metrics were derived for AI, PI and Sham was: 720 ± 113; 721 ± 125 and 738 ± 130. The mean heart rate for AI, PI and Sham during LIFU/CHEPS administration was: 72.4 bpm ± 11.5 bpm; 71.3 bpm ± 11.4 bpm and 73.3 bpm ± 11.8 bpm. No significant differences were found (p > 0.05) (**Figure 5A**).

**Figure 5.**
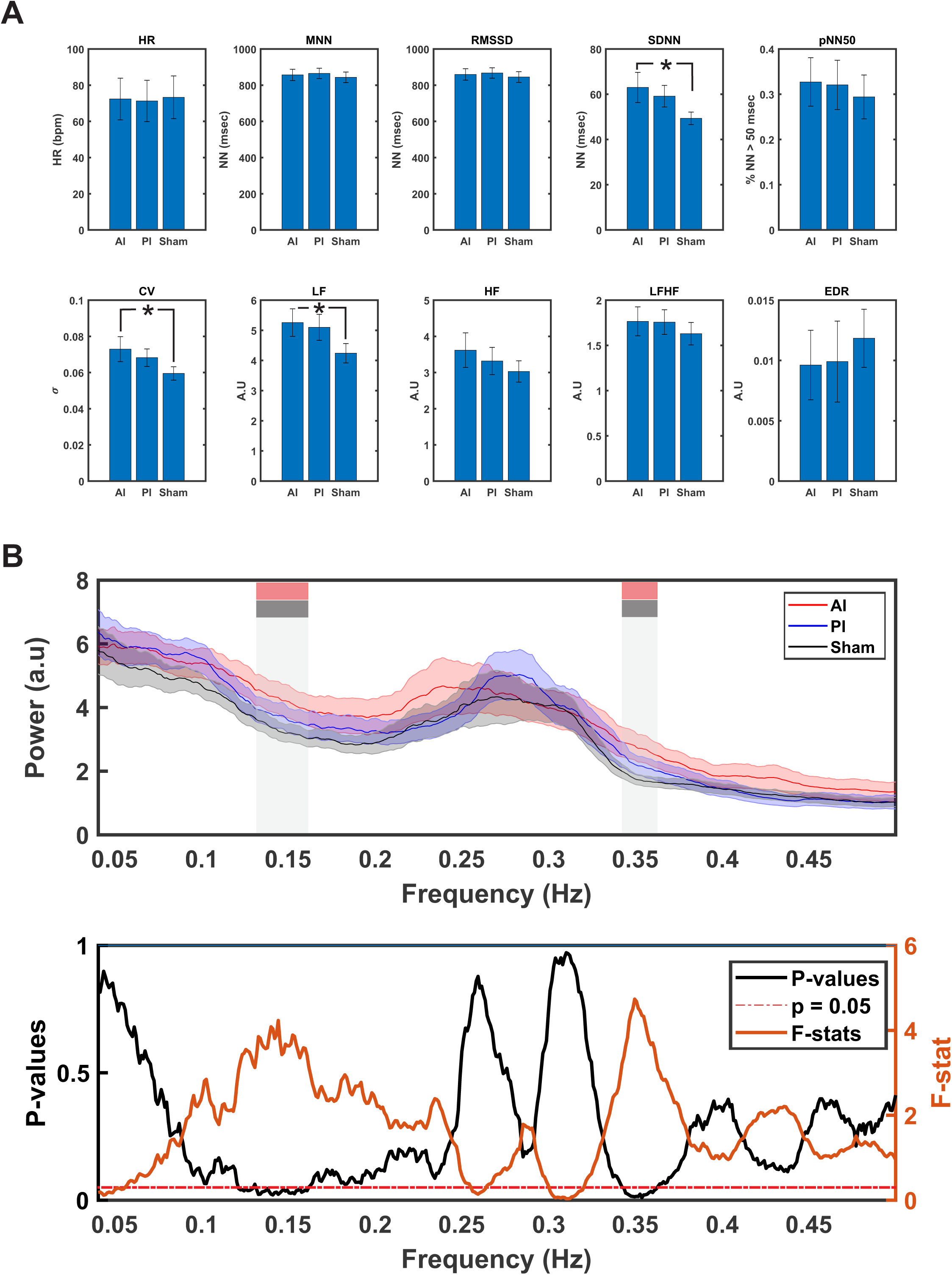
Effect of LIFU on autonomic measures. **A.** Group (N = 23) autonomic measures. Bars are mean ± SEM. * denote significant difference (p< 0.05) between AI and Sham only. HR = heart rate; MNN = mean NN interval; RMSDD = root mean square difference; SDNN = standard deviation of NN peaks; pNN50 = proportion of NN peaks < 50 msec; CV = coefficient of variation; LF = low frequency power; HF = high frequency power; LFHF = low/high frequency power ratio; EDR = electrodermal response. **B.** (Top) Group (N = 23) mean ± SEM heart rate power spectra. Vertical gray bars denote timing of significant differences in magnitude of power spectra from permutation statistics (5000 randomizations, p < 0.05) (shown below). Colored boxes on gray bars denote conditions that were statistically significant (AI (red) & Sham (black)).

#### Heart rate variability

Time domain parameters of interest included mean of the RR intervals (MNN), root mean square difference of successive RR intervals (RMSSD), standard deviation of the RR intervals (SDNN) and the coefficient of variation (CV = σ/µ). Frequency domain parameters of interest included low-frequency (LF; 0.04 – 0.15 Hz), high frequency (HF; 0.151 – 0.4 Hz) power and the LF/HF ratio.

#### MNN

The mean MNN for AI, PI and Sham were: 857 ± 151, 865 ± 138 and 843 ± 141 milliseconds respectively. No significant differences were found; F(2,44) = 0.78, p = 0.46 (**Figure 5A**).

#### RMSSD

The mean RMSSD for AI, PI and Sham were: 860 ± 152, 868 ± 138 and 845 ± 141 milliseconds respectively. No significant differences were found; F(2,44) = 0.82, p = 0.45 (**Figure 5A**).

#### SDNN

The mean SDNN for AI, PI and Sham were: 63 ± 32, 59 ± 23 and 49 ± 13 milliseconds respectively. The one-way repeated measures ANOVA revealed a significant difference: F(2,44) = 3.75, p = 0.03. Post-hoc Tukey tests revealed the main effect to be the result of a significant difference between Sham and AI (p < 0.05) (**Figure 5A**).

#### pNN50

The mean ± SD for AI, PI and Sham were: 0.33 ± 0.25, 0.32 ± 0.26 and 0.29 ± 0.23. The one-way repeated measures ANOVA revealed no significant main effect: F(2,44) = 0.50, p = 0.61 (**Figure 5A**).

#### Coefficient of Variation (CV)

The mean CV for AI, PI and Sham were: 0.073 ± 0.03, 0.068 ± 0.02 and 0.059 ± 0.01. The one-way repeated measures ANOVA revealed a significant difference: F(2,44) = 3.42, p = 0.041. Post-hoc Tukey tests revealed the main effect to be the result of a significant difference between Sham and AI (p < 0.05) (**Figure 5A**).

#### LF

The mean power for AI, PI and Sham were: 0.0053 ± 0.002, 0.0051 ± 0.002 and 0.0042 ± 0.0015. The one-way repeated measures ANOVA revealed a significant difference: F(2,44) = 3.98, p = 0.025. Post-hoc Tukey tests revealed the main effect to be the result of a significant difference between Sham and AI (p < 0.05) (**Figure 5A**).

#### HF

The mean power for AI, PI and Sham were: 0.0036 ± 0.002, 0.0033 ± 0.001 and 0.003 ± 0.001. No significant differences were found: F(2,44) = 1.68, p = 0.2 (**Figure 5A**).

#### LFHF

The mean power ratio for AI, PI and Sham were: 1.77 ± 0.77, 1.76 ± 0.65 and 1.63 ± 0.59. No significant differences were found: F(2,44) = 0.81, p = 0.45 (**Figure 5A**).

#### EDR

The mean peak-to-peak EDR conductive response in microseiverts for AI, PI and Sham were: 0.01 ± 0.01, 0.01 ± 0.02 and 0.011 ± 0.01. No significance differences were found: F(2,44) = 0.33, p = 0.72 (**Figure 5A**).

We further investigated the frequency domain HRV data to determine if the LF differences were due to a specific frequency within the band. For this analysis we ran non-parametric permutation statistics (p < 0.05, 5000 randomizations) across each of the frequencies 0.04 to 0.4 with a 0.01 Hz resolution. In addition, a temporal cluster threshold of 10 consecutive points (0.1 Hz) was required to satisfy significance. This analysis revealed two significant clusters: One cluster from 0.131 – 0.16 Hz and another cluster at 0.342 – 0.362 (p < 0.05). Post-hoc analysis of these clusters found that each was driven by a difference between Sham and AI (p < 0.05) (see **Figure 5B**).

### Supplemental Data

#### Estimated in vivo pressure

All participants received 400 kPa to the temporal window. The estimated *in vivo* mean ± SD pressure at the insula combining both AI and PI sonications was: 221.4 ± 93.9 kPa with a range of 72.7 to 460.3 kPa for an overall attenuation of 45%. The mean ± SD for AI was 179.3 ± 75.1 kPa with a range of 72.7 to 350.7 kPa. Mean attenuation for AI was 55%. The mean ± SD for PI was 263.5 ± 93.4 kPa with a range of 114.6 to 460.3 kPa. The mean attenuation for PI was 34%. A paired t-test revealed a significant difference: t(22) = −3.24, p = 0.0038 such that the estimated pressure in the head for the AI condition was significantly lower than for PI (**Figure S1**).

#### Skull density ratio and thickness

The mean ± SD skull thickness at the left temporal window was 3.55 ± 0.68 mm with a range of 2.44 to 4.99 mm. The average skull density ratio (SDR) was 0.78 ± 0.09 with a range of 0.59 to 0.95 (**Figure S1**).

In an effort to help explain the variance in the estimated *in vivo* pressure, we performed separate Pearson’s correlations between pressure and SDR and thickness. Using both AI and PI data points together there was no linear relationship between estimated *in vivo* pressure at the insula target with SDR (r = 0.1, p = 0.49) or skull thickness (r = −0.17, p = 0.26).

There is currently no data on the dosing of LIFU in human applications. In an effort to relate estimated in vivo to effect size we performed separate Pearson’s correlations between pressure and percent perceived pain change score and percent change peak-to-peak CHEP amplitude as compared to Sham. Using both AI and PI data points together there was no linear relationship between estimated pressure at the insula with perceived pain ratings (r = −0.04, p = 0.8) or with CHEP N1/P1 peak-to-peak amplitude (r = −0.05, p = 0.72) (**Figure S1**).

#### Targeting Error

The mean ± SD targeting error across all participants for conditions and all trials was 1.19 ± 0.35 mm. We further broke this down by participant and by condition. The mean targeting error for AI, PI and Sham conditions were 1.21 ± 0.66; 1.25 ± 0.65 and 1.16 ± 0.64. The range of mean error across subjects was 0.6 – 2.1 mm (see **Figure S2**).

#### Questionnaires

We queried participants using the Beck Depression Inventory, State and Trait Anxiety Inventory and the Pain Catastrophizing Scale. The mean ± SD for BDI was 4.5 ± 3.9 (range 1 – 18). The mean ± SD for the STAI-T was 37.3 ± 7.6 (range 27 – 57). The mean ± for the PCS was: 4.9 ± 4.2 (range 0 – 13).

#### Report of symptoms

We queried participants both before and 30 minutes after each formal testing session using report of symptoms questionnaire previously employed in our LIFU studies[82] on adverse events and symptom severity. If a symptom or AE was present the participant was requested to rate the severity as either mild, moderate or severe. No severe AE or symptoms were reported for any condition (AI, PI, Sham) either before or after the intervention. Sleepiness was the symptom that was most frequently reported but was not specific to an intervention or time period before or after intervention. Group cumulative symptom reports as well as individual subject differences between pre and post intervention for each condition (AI, PI, Sham) is presented in **Figure S2**.

#### Acoustic masking

We queried participants 30 minutes after the conclusion of formal testing of each session whether they could hear or feel the LIFU and whether they believed they had received a LIFU intervention using 0 – 6 Likert scales. The mean ± SD for the AI, PI and Sham sessions for the question ‘I could hear the LIFU stimulation’ were: 2.4 ± 2.1, 1.7 ± 1.9 and 1.7 ± 1.9 equating to a ‘Disagree’ rating. The Kruskalwallis test revealed no statistically significant differences between groups: (df = 2,66), Chi-sq = 1.83, p = 0.39. Of the N = 23 only 1 person strongly agreed (6 on 0 – 6 scale) that they could hear the LIFU during one of the active conditions (PI) however they also strongly agreed they could hear it on the Sham condition too (see **Figure S2**). For the question ‘I could feel the LIFU stimulation, the mean ± SD for the AI, PI and Sham sessions were: 0.9 ± 1.4, 1.0 ± 1.4 and 1.0 ± 1.7 equating to a response of ‘Disagree’. The Kruskalwallis test revealed no statistically significant differences between groups: df = (2,66), Chi-square = 0.18, p = 0.92. There were only 3 total responses for the Agree and Strongly Agree responses and all were made by the same participants for all conditions (including Sham) (see **Figure S2**). For the question, ‘I believe I experienced LIFU stimulation’ the mean ± SD for the AI, PI and Sham conditions were: 3.2 ± 1.8, 3.6 ± 1.2 and 3.1 ± 1.5 equating to a response of ‘Neutral’ suggesting that the participants did not know either way. The Kruskalwallis test revealed no significant difference between groups: df = (2,66), Chi-square = 1.14, p = 0.56 (**Figure S2**).

## DISCUSSION

This is the first study in humans (or otherwise) to test the effect of LIFU to sub-regions of the insula for effects on pain and autonomic function. We delivered single-element 500 kHz LIFU to either the AI or PI as compared to an in-active Sham for effects on perceived pain, EEG metrics from contact heat and indices of autonomic reactivity including heart-rate variability, electrodermal response and blood pressure. Only LIFU to PI significantly reduced perceived pain ratings compared to Sham and only LIFU to AI affected autonomic measures of heart-rate variability. However, LIFU to both AI and PI affected EEG activity. LIFU to PI exclusively affected earlier EEG activity from 290 – 313 msec whereas LIFU to AI exclusively affected later EEG activity from 485 – 504 msec.

Reduction in perceived pain may be the result of either decreased bottom-up coding of stimulus attributes or from top-down salience encoding mechanisms. The spinal-medullary-spinal pain inhibitory system[95] is an endogenous pathway of descending pain control that can serve to gate or inhibit incoming pain signals to cortical areas like the posterior insula. Top-down mechanisms involve eloquent brain regions such as the dorsal anterior cingulate cortex (dACC) and anterior insula that have strong connections with periaqueductal gray and have been implicated in the descending control of pain[96,97]. Another option to explain natural reduction in perceived pain is a reduction in stimulus saliency or novelty[98,99]. LIFU to either the AI or PI did not remove this inhibition over the course of the 40 trials but LIFU to PI was significantly lower from bins 5-7 suggesting an effect on rate of inhibition. Whether this is a result of pain inhibition mechanisms or stimulus saliency is debatable. The precise role of insula in nociception, pain and the overall pain experience is incompletely understood. While insula activity is a common finding in human studies employing noxious or painful stimuli[31,41] and has been identified as part of a dynamic pain connectome[38], this activity may not specifically index pain *per se* but rather reflect attention to or salience of the stimlus[55,100] as the dorsal anterior insula is a critical hub in the so-called salience network[56,101]. Indeed, the debate of what insula activity specifically represents during human studies of pain is charged and highly active[34,102,103]. Regardless, it is clear that spinothalamic afferents from lamina I of the spinal cord that carry A-delta and C-fiber mediated nociceptive information primarily terminate in the posterior insula[33,47,59] and that the posterior insula codes physical stimulus attributes including intensity, modality and somatotopy[33,59]. The PI is highly interconnected with the AI and it is postulated that a hierarchical serial processing chain exists in the insula for noxious stimuli whereby stimulus attributes are passed from the PI to AI where salience, emotion and expectation are integrated to produce to overall subjective percept of pain[35,44,104]. While LIFU to AI and PI had similar mean effects on perceived pain, it is inconclusive from this study if LIFU to PI worked by reducing stimulus intensity coding whereas LIFU to AI worked by reducing stimulus saliency. Evidence for these particular roles of AI and PI to pain processing was examined in an interesting study by Lutz et al. (2013)[105] where highly practiced meditators reported no change in the intensity of a painful stimulus but did report reduced unpleasantness and that these differences were indexed by activity in the anterior insula. LIFU provides another potential means of disentangling issues of pain processing related to different areas of the insula and salience.

The CHEP has a similar waveform morphology to the laser evoked potential (LEP) in that contact heat produces a large (~ 40 uV) negative/positive complex though the latency looks to be longer than an LEP[89]. Nevertheless, CHEP are well studied[88,106,106], have been source localized to the insula[107] and activate similar brain regions to lasers as measured using fMRI[108]. Given the strong similarities of CHEP with LEP, it is useful to understand our results from some of the LEP literature. Using intracerebral recordings, Frot et al.[35] evaluated the contribution of different insula gyri to the amplitude of the LEP and reported amplitudes to be largest and earlier for posterior insula suggesting a greater contribution of posterior insula to the generation of the of the LEP as compared to the anterior insula though phase reversals were identified suggesting distinct generators in anterior insula as well. LIFU provides a potential non-invasive method to determine the causal contribution of different insula sub-regions to the CHEP. Anatomical tracing work by Craig[33,59] and deep brain recordings by Frot et al.[35] suggest a serial hierarchical model of pain processing within the insula which progresses in space and time from PI (stimulus attribute coding (modality, intensity, somatotopy)) to AI (salience, emotion).

Given this, it is reasonable to assume that the N1 (which has a shorter latency) indexes an earlier (and presumably more posterior) window of processing as compared to the P1 (which is later and hence may index anterior insula activity). Under this assumption, one would hypothesize that LIFU to the PI would preferentially affect the N1 whereas LIFU to AI would preferentially affect the P1. While the peak-to-peak analysis demonstrated LIFU to AI and PI both attenuated the peak-to-peak amplitude of the CHEP driven by exclusive effects on the P1, the permutation analysis did indeed reveal specific effects for LIFU to PI to exclusively affect the CHEP trace at earlier timings around the N1 (but not at the N1 peak timing) whereas LIFU to AI exclusively affected the CHEP trace coincident with the timing of the P1 peak. A final finding from the permutation analysis revealed a main effect at 824-903 msec driven by significant differences between both AI and PI as compared to Sham such that LIFU to AI and PI both reduced the amplitude at this time point. The underlying cause for both AI and PI effects at the 824-903 msec time point could conceivably index later integrative processing combining bottom-up signals with top-down expectation or predictions involving both the posterior and anterior insula as set out in the Embodied Predictive Interoception Coding model[109]. While this is purely speculative, it is notable that specific PI and AI effects were found early but both as compared to Sham much later. Another interpretation of evoked potentials is that they do not represent a serial processing stream but rather represent phase locking of brain oscillations within specific frequency bands[110,111]. This interpretation has been posited for somatosensory evoked potentials[112]. It is clear from our time-frequency data that the timing of the CHEP minima/maxima is associated with high time-locked power in the 2 – 13 Hz range and that inhibition of CHEP amplitude is reflected as a decrease in power of these frequencies that is clearly time-locked around the timing of the CHEP. Interestingly, effects on EEG power spectra were largely the result of LIFU to the anterior insula and not the posterior insula. We found significant power reductions in delta, theta and alpha power concentrated in the time window around the generation of the CHEP (200 – 600 msec). Delta power (in waking) is believed to be generated by anterior medial frontal cortex including cingulate regions, insula, nucleus accumbens and ventral tegmental area[113] – areas traditionally regarded as being involved in pleasure, reward and addiction though there is substantial research demonstrating the role of nucleus accumbens in the mediation of pain[114]. Of considerable interest is the link between delta power and homeostatic processes[115] including heart rate variability that showed an inverse relationship[116,117] similar to our results where LIFU to AI reduced delta power but increased SDNN. The effects in the other frequency ranges may index pain processing. According to Ploner et al. (2017)[118] the sending of feedforward information is associated with gamma frequency whereas if pain is driven by contextual top-down process is associated with alpha/beta frequencies. Here, differences in power were largely driven by LIFU to the AI and were reflected in decreased power in mainly theta and alpha ranges. The role of theta frequencies in phasic or acute pain is not well understood however, chronic pain appears to be associated with abnormal oscillations at theta frequencies and linked to abnormal contextual feedback processes (AI and salience and context, expectation, motivation etc.) As such, LIFU to AI particularly reduced the power of these frequency band which may make it a candidate intervention for chronic pain as opposed to the PI that did not demonstrate this effect.

### Autonomic results

We found effects of LIFU to AI on autonomic metrics including SDNN, CV, mean low-frequency power but also in two distinct frequency bands (0.131 – 0.16 Hz and 0.342 – 0.362 Hz) that span both the formal LF and HF frequency ranges. That only LIFU to AI showed effects would concur with structural anatomy as the anterior insula (among other brain areas including anterior cingulate cortex, central nucleus of the amygdala and hypothalamic nuclei) is one origin of efferent fibers that project to medullary and spinal nuclei controlling cardiac function[65]. This is opposed to the posterior insula that looks to predominantly receive cardiac afferents via thalamic relay nuclei[65]. Chouchou et al (2019)[68] performed functional mapping of autonomic cardiac responses using direct electric stimulation of the insula in awake humans and found, in general, that stimulation of the anterior insula produced more bradycardia concomitant with an increase in parasympathetic tone as indexed by the increase in power of HF whereas stimulation of the PI predominantly produced tachycardia accompanied by an increase in LF/HF ratio suggesting an increase in sympathetic tone. Our results support Craig’s model (based upon structural anatomy) and are encouraging as LIFU to AI increased SDNN (an increase in vagal tone) that is generally regarded as positive for overall health, whereas decreases in HRV are associated with poorer health and neuropsychological disease[119,120].

### Safety

LIFU for human neuromodulation generally follows the IEC standard 60601 part 1[121] or FDA guidelines for diagnostic imaging[122] and looks to have a favorable safety profile[8,82]. We previously reported on adverse events in N = 64 healthy volunteer participants that received LIFU under several different protocols to different brain areas at different intensities and found no serious events and a safety profile comparable to other forms of non-invasive neuromodulation[82]. Here, no serious adverse events were reported with the most common symptom being sleepiness though this occurred at similar rates for all testing sessions including sham suggesting it was not specific to the intervention.

## CONCLUSIONS & FUTURE WORK

500 kHz single-element LIFU is an effective non-invasive means to transiently modulate activity in specific insular sub-regions that also affects behavior. Future work will look to establish minimal effective dose as well as the longevity of the effect. These will be important considerations for the successful translation of LIFU as a potential therapeutic for clinical indications such as chronic pain.

## ACKNOWLEDGMENTS

This work was funded in part by grants awarded to WL from the Seale Innovation Fund and the National Institute for Health (NIH) 1R21AT012247-01. The authors would like to thank Jessica Florig for help with data collection and analysis.

## DATA AVAILABILITY STATEMENT

The authors comply with PLOS’ data policy. Data and source code is available upon reasonable request.

---

**Table S1.** Insular Targeting.

|  |  | MNI Coordinates |  |  |  |  |  | Target Depth (mm) |  |
| --- | --- | --- | --- | --- | --- | --- | --- | --- | --- |
|  |  | X | Y | Z | X | Y | Z | AI | PI |
| **Male** | S1 | -31 | 22 | -3 | -39 | -6 | 4 | 40.8 | 42.4 |
|  | S2 | -37 | 18 | -4 | -37 | -4 | 0 | 30.4 | 35.8 |
|  | S3 | -33 | 21 | -6 | -36 | -10 | -1 | 37.7 | 45 |
|  | S4 | -32 | 26 | -5 | -35 | -5 | 4 | 34.2 | 40.8 |
|  | S5 | -38 | 21 | 1 | -39 | -3 | 2 | 30.4 | 35.8 |
|  | S6 | -33 | 14 | 0 | -34 | -12 | 4 | 35.8 | 42 |
|  | S7 | -33 | 24 | -3 | -36 | -11 | 2 | 35.8 | 40.7 |
| **Female** | S8 | -31 | 16 | -1 | -34 | -14 | 1 | 36.3 | 41.9 |
|  | S9 | -35 | 13 | -5 | -34 | -13 | 1 | 33.2 | 40.4 |
|  | S10 | -33 | 23 | -1 | -35 | -4 | -1 | 34.5 | 38.9 |
|  | S11 | -37 | 21 | -6 | -39 | -6 | -4 | 33.6 | 36.9 |
|  | S12 | -36 | 17 | -1 | -36 | -4 | 2 | 30.2 | 38 |
|  | S13 | -33 | 21 | -3 | -34 | -4 | 1 | 31.2 | 37.3 |
|  | S14 | -35 | 19 | -8 | -36 | -5 | 5 | 30.9 | 37.7 |
|  | S15 | -33 | 19 | -4 | -36 | -10 | -1 | 33.4 | 38.1 |
|  | S16 | -35 | 20 | -2 | -37 | -9 | -1 | 34.7 | 39.7 |
|  | S17 | -29 | 22 | -1 | -33 | -5 | -3 | 39.5 | 44.1 |
|  | S18 | -34 | 19 | -5 | -33 | -10 | 3 | 36.2 | 41.2 |
|  | S19 | -34 | 18 | -4 | -35 | -5 | 2 | 34.9 | 40.3 |
|  | S20 | -34 | 21 | -4 | -35 | -7 | 0 | 34.3 | 38.1 |
|  | S21 | -31 | 20 | -4 | -37 | -8 | 4 | 34 | 37.2 |
|  | S22 | -35 | 17 | -2 | -35 | -7 | 1 | 32.3 | 38.8 |
|  | S23 | -32 | 24 | -3 | -36 | -7 | -2 | 37.5 | 42.4 |

**Supplementary Figure 1.**
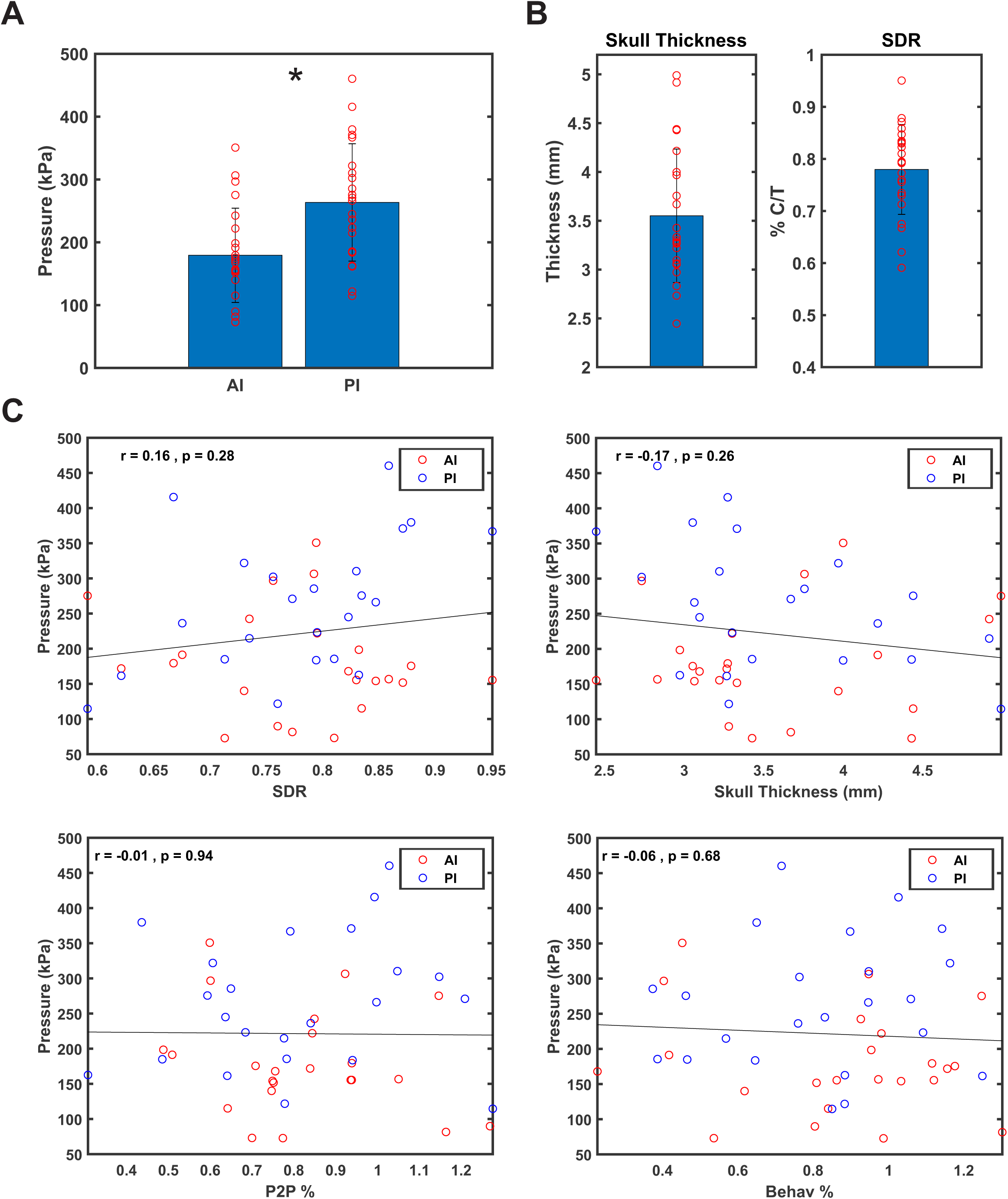
LIFU pressure and skull effects. **A.** Group (N = 23) estimated *in vivo* pressure from acoustic modelling at the anterior insula (AI) and posterior insula (PI) targets. Bars are mean ± SEM. Individual participant data is shown in red. * denotes significant difference (p< 0.05). **B.** (Left) Group (N = 23) skull thickness. Bars are mean ± SEM. Individual participant data is shown in red. (Right) Group (N = 23) skull-density ratio (SDR). Bars are mean ± SEM. Individual participant data is shown in red. **C.** Group (N = 23) correlations of estimated in vivo pressure at both the anterior insula (AI) and posterior insula (PI) targets with SDR (top left), skull thickness (top right), peak-to-peak CHEP amplitude percent change from Sham (P2P%) and perceived pain rating percent change from Sham (Behav%).

**Supplementary Figure 2.**
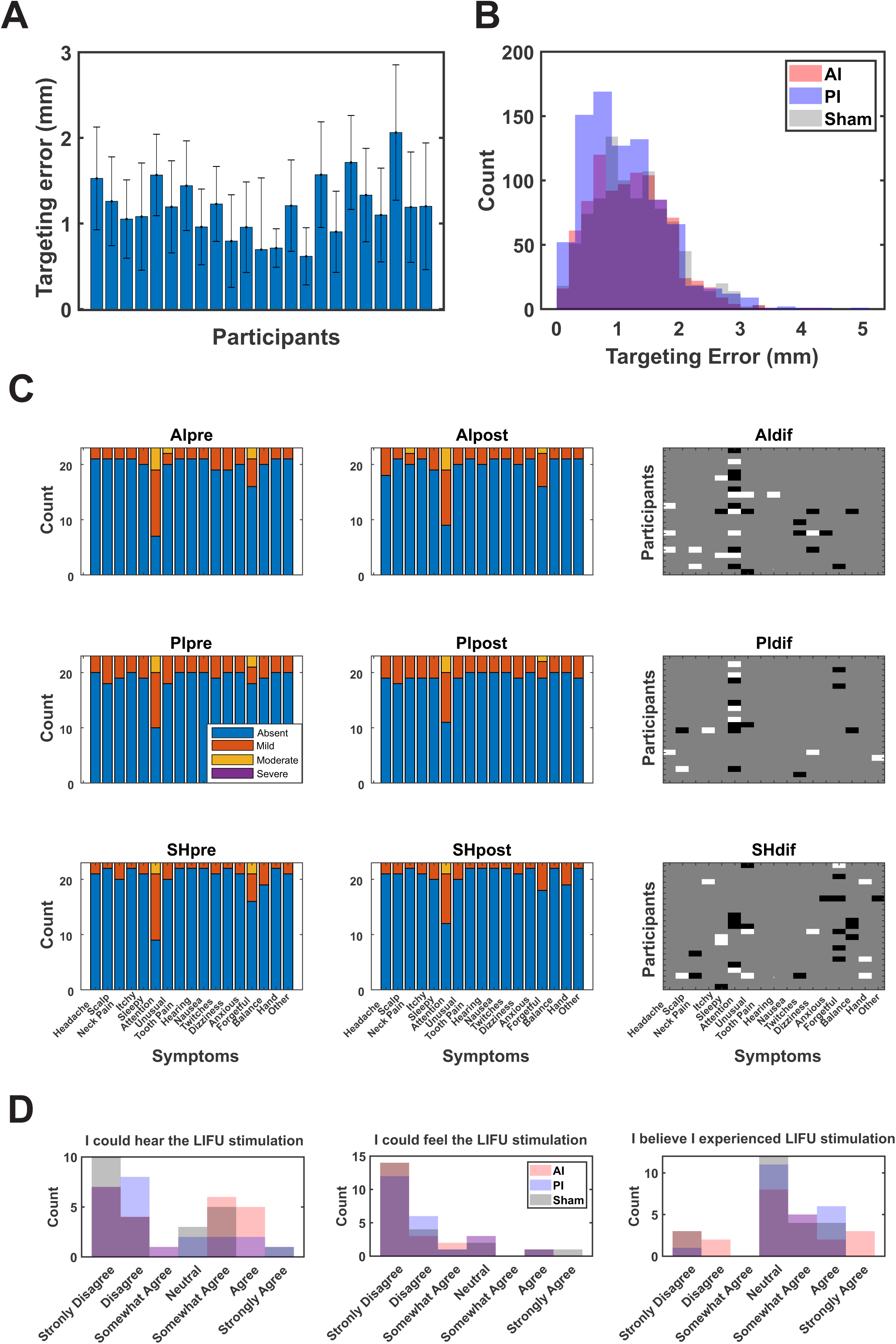
Targeting, Safety and Acoustic masking. **A.** Individual participants targeting error (mean ± SEM) of the LIFU transducer on the scalp target pooled across both AI and PI targets. **B.** Histogram of targeting error of transducer placement on the scalp for all targets from all participants for each LIFU condition. There were no significant differences between conditions. **C.** Group (N = 23) adverse event data recorded before LIFU (left column) and 30 minutes after LIFU (middle column) for each condition. Top row = anterior insula (AI); middle row = posterior insula (PI); bottom row = Sham. Right most column shows individual participant differences in reporting before/after LIFU for each respective condition. Black = 1pt down; White = 1pt up. No participant reported an increase or decrease in severity of a reported symptom by more than 1pt. **D.** Group (N =23) histograms from the acoustic mask evaluation separated by condition.

## REFERENCES

1. Deng Z-D, Lisanby SH, Peterchev AV. Electric field depth–focality tradeoff in transcranial magnetic stimulation: simulation comparison of 50 coil designs. Brain stimulation. 2013;6: 1–13.

2. Gomez-Tames J, Hamasaka A, Hirata A, Laakso I, Lu M, Ueno S. Group-level analysis of induced electric field in deep brain regions by different TMS coils. Phys Med Biol. 2020;65: 025007. doi:10.1088/1361-6560/ab5e4a

3. Spagnolo PA, Wang H, Srivanitchapoom P, Schwandt M, Heilig M, Hallett M. Lack of Target Engagement Following Low-Frequency Deep Transcranial Magnetic Stimulation of the Anterior Insula. Neuromodulation: Technology at the Neural Interface. 2019;22: 877– 883. doi:10.1111/ner.12875

4. Coll M-P, Penton T, Hobson H. Important methodological issues regarding the use of transcranial magnetic stimulation to investigate interoceptive processing: a Comment on Pollatos et al. (2016). Philosophical Transactions of the Royal Society B: Biological Sciences. 2017;372: 20160506. doi:10.1098/rstb.2016.0506

5. Legon W, Sato TF, Opitz A, Mueller J, Barbour A, Williams A, et al. Transcranial focused ultrasound modulates the activity of primary somatosensory cortex in humans. Nature neuroscience. 2014;17: 322–329.

6. Legon W, Ai L, Bansal P, Mueller JK. Neuromodulation with single-element transcranial focused ultrasound in human thalamus. Human brain mapping. 2018;39: 1995–2006.

7. Darmani G, Bergmann T, Pauly KB, Caskey C, de Lecea L, Fomenko A, et al. Non-invasive transcranial ultrasound stimulation for neuromodulation. Clinical Neurophysiology. 2021.

8. Blackmore J, Shrivastava S, Sallet J, Butler CR, Cleveland RO. Ultrasound Neuromodulation: A Review of Results, Mechanisms and Safety. Ultrasound in Medicine & Biology. 2019;45: 1509–1536. doi:10.1016/j.ultrasmedbio.2018.12.015

9. Bystritsky A, Korb AS. A Review of Low-Intensity Transcranial Focused Ultrasound for Clinical Applications. Curr Behav Neurosci Rep. 2015;2: 60–66. doi:10.1007/s40473-015-0039-0

10. Sassaroli E, Vykhodtseva N. Acoustic neuromodulation from a basic science prospective. J Ther Ultrasound. 2016;4: 17. doi:10.1186/s40349-016-0061-z

11. Naor O, Krupa S, Shoham S. Ultrasonic neuromodulation. J Neural Eng. 2016;13: 031003. doi:10.1088/1741-2560/13/3/031003

12. Tufail Y, Matyushov A, Baldwin N, Tauchmann ML, Georges J, Yoshihiro A, et al. Transcranial Pulsed Ultrasound Stimulates Intact Brain Circuits. Neuron. 2010;66: 681–694. doi:10.1016/j.neuron.2010.05.008

13. King RL, Brown JR, Newsome WT, Pauly KB. Effective Parameters for Ultrasound-Induced In Vivo Neurostimulation. Ultrasound in Medicine & Biology. 2013;39: 312–331. doi:10.1016/j.ultrasmedbio.2012.09.009

14. Kim H, Chiu A, Lee SD, Fischer K, Yoo S-S. Focused Ultrasound-mediated Non-invasive Brain Stimulation: Examination of Sonication Parameters. Brain Stimulation. 2014;7: 748–756. doi:10.1016/j.brs.2014.06.011

15. Yu K, Niu X, Krook-Magnuson E, He B. Intrinsic functional neuron-type selectivity of transcranial focused ultrasound neuromodulation. Nat Commun. 2021;12: 2519. doi:10.1038/s41467-021-22743-7

16. Yoo S-S, Bystritsky A, Lee J-H, Zhang Y, Fischer K, Min B-K, et al. Focused ultrasound modulates region-specific brain activity. NeuroImage. 2011;56: 1267–1275. doi:10.1016/j.neuroimage.2011.02.058

17. Lee W, Lee SD, Park MY, Foley L, Purcell-Estabrook E, Kim H, et al. Image-Guided Focused Ultrasound-Mediated Regional Brain Stimulation in Sheep. Ultrasound in Medicine & Biology. 2016;42: 459–470. doi:10.1016/j.ultrasmedbio.2015.10.001

18. Dallapiazza RF, Timbie KF, Holmberg S, Gatesman J, Lopes MB, Price RJ, et al. Noninvasive neuromodulation and thalamic mapping with low-intensity focused ultrasound. Journal of Neurosurgery. 2018;128: 875–884. doi:10.3171/2016.11.JNS16976

19. Folloni D, Verhagen L, Mars RB, Fouragnan E, Constans C, Aubry J-F, et al. Manipulation of Subcortical and Deep Cortical Activity in the Primate Brain Using Transcranial Focused Ultrasound Stimulation. Neuron. 2019;101: 1109–1116.e5. doi:10.1016/j.neuron.2019.01.019

20. Kubanek J, Brown J, Ye P, Pauly KB, Moore T, Newsome W. Remote, brain region– specific control of choice behavior with ultrasonic waves. Sci Adv. 2020;6: eaaz4193. doi:10.1126/sciadv.aaz4193

21. Deffieux T, Younan Y, Wattiez N, Tanter M, Pouget P, Aubry J-F. Low-Intensity Focused Ultrasound Modulates Monkey Visuomotor Behavior. Current Biology. 2013;23: 2430–2433. doi:10.1016/j.cub.2013.10.029

22. Wattiez N, Constans C, Deffieux T, Daye PM, Tanter M, Aubry J-F, et al. Transcranial ultrasonic stimulation modulates single-neuron discharge in macaques performing an antisaccade task. Brain Stimulation. 2017;10: 1024–1031. doi:10.1016/j.brs.2017.07.007

23. Yang P-F, Phipps MA, Newton AT, Chaplin V, Gore JC, Caskey CF, et al. Neuromodulation of sensory networks in monkey brain by focused ultrasound with MRI guidance and detection. Sci Rep. 2018;8: 7993. doi:10.1038/s41598-018-26287-7

24. Legon W, Bansal P, Tyshynsky R, Ai L, Mueller JK. Transcranial focused ultrasound neuromodulation of the human primary motor cortex. Scientific reports. 2018;8: 1–14.

25. Lee W, Kim H-C, Jung Y, Chung YA, Song I-U, Lee J-H, et al. Transcranial focused ultrasound stimulation of human primary visual cortex. Sci Rep. 2016;6: 34026. doi:10.1038/srep34026

26. Fomenko A, Chen K-HS, Nankoo J-F, Saravanamuttu J, Wang Y, El-Baba M, et al. Systematic examination of low-intensity ultrasound parameters on human motor cortex excitability and behavior. eLife. 2020;9: e54497. doi:10.7554/eLife.54497

27. Mueller J, Legon W, Opitz A, Sato TF, Tyler WJ. Transcranial focused ultrasound modulates intrinsic and evoked EEG dynamics. Brain stimulation. 2014;7: 900–908.

28. Ai L, Mueller JK, Grant A, Eryaman Y, Legon W. Transcranial focused ultrasound for BOLD fMRI signal modulation in humans. IEEE; 2016. pp. 1758–1761.

29. Ai L, Bansal P, Mueller JK, Legon W. Effects of transcranial focused ultrasound on human primary motor cortex using 7T fMRI: a pilot study. BMC neuroscience. 2018;19: 1–10.

30. Bergeron D, Obaid S, Fournier-Gosselin M-P, Bouthillier A, Nguyen DK. Deep Brain Stimulation of the Posterior Insula in Chronic Pain: A Theoretical Framework. Brain Sciences. 2021;11: 639. doi:10.3390/brainsci11050639

31. Brooks JCW, Tracey I. The insula: A multidimensional integration site for pain. Pain. 2007;128: 1–2. doi:10.1016/j.pain.2006.12.025

32. Lu C, Yang T, Zhao H, Zhang M, Meng F, Fu H, et al. Insular Cortex is Critical for the Perception, Modulation, and Chronification of Pain. Neurosci Bull. 2016;32: 191–201. doi:10.1007/s12264-016-0016-y

33. Craig A. Pain mechanisms: labeled lines versus convergence in central processing. Annual review of neuroscience. 2003;26: 1–30.

34. Segerdahl AR, Mezue M, Okell TW, Farrar JT, Tracey I. The dorsal posterior insula subserves a fundamental role in human pain. Nat Neurosci. 2015;18: 499–500. doi:10.1038/nn.3969

35. Frot M, Faillenot I, Mauguière F. Processing of nociceptive input from posterior to anterior insula in humans: Nociceptive Input Processing in the Insula. Hum Brain Mapp. 2014;35: 5486–5499. doi:10.1002/hbm.22565

36. Baliki MN, Apkarian AV. Nociception, Pain, Negative Moods, and Behavior Selection. Neuron. 2015;87: 474–491. doi:10.1016/j.neuron.2015.06.005

37. Davis KD, Moayedi M. Central Mechanisms of Pain Revealed Through Functional and Structural MRI. J Neuroimmune Pharmacol. 2013;8: 518–534. doi:10.1007/s11481-012-9386-8

38. Kucyi A, Davis KD. The dynamic pain connectome. Trends in Neurosciences. 2015;38: 86–95. doi:10.1016/j.tins.2014.11.006

39. Garcia-Larrea L, Peyron R. Pain matrices and neuropathic pain matrices: A review. Pain. 2013;154: S29–S43. doi:10.1016/j.pain.2013.09.001

40. Garcia-Larrea L, Frot M, Valeriani M. Brain generators of laser-evoked potentials: from dipoles to functional significance. Neurophysiologie Clinique/Clinical Neurophysiology. 2003;33: 279–292. doi:10.1016/j.neucli.2003.10.008

41. Wager TD, Atlas LY, Lindquist MA, Roy M, Woo C-W, Kross E. An fMRI-Based Neurologic Signature of Physical Pain. N Engl J Med. 2013;368: 1388–1397. doi:10.1056/NEJMoa1204471

42. Faillenot I, Heckemann RA, Frot M, Hammers A. Macroanatomy and 3D probabilistic atlas of the human insula. NeuroImage. 2017;150: 88–98. doi:10.1016/j.neuroimage.2017.01.073

43. Cauda F, D’Agata F, Sacco K, Duca S, Geminiani G, Vercelli A. Functional connectivity of the insula in the resting brain. NeuroImage. 2011;55: 8–23. doi:10.1016/j.neuroimage.2010.11.049

44. Craig AD, Craig A. How do you feel--now? The anterior insula and human awareness. Nature reviews neuroscience. 2009;10.

45. Kelly C, Toro R, Di Martino A, Cox CL, Bellec P, Castellanos FX, et al. A convergent functional architecture of the insula emerges across imaging modalities. NeuroImage. 2012;61: 1129–1142. doi:10.1016/j.neuroimage.2012.03.021

46. Kurth F, Zilles K, Fox PT, Laird AR, Eickhoff SB. A link between the systems: functional differentiation and integration within the human insula revealed by meta-analysis. Brain Struct Funct. 2010;214: 519–534. doi:10.1007/s00429-010-0255-z

47. Dum RP, Levinthal DJ, Strick PL. The Spinothalamic System Targets Motor and Sensory Areas in the Cerebral Cortex of Monkeys. Journal of Neuroscience. 2009;29: 14223–14235. doi:10.1523/JNEUROSCI.3398-09.2009

48. Frot M, Magnin M, Mauguière F, Garcia-Larrea L. Human SII and Posterior Insula Differently Encode Thermal Laser Stimuli. Cerebral Cortex. 2007;17: 610–620. doi:10.1093/cercor/bhk007

49. Baumgärtner U, Iannetti GD, Zambreanu L, Stoeter P, Treede R-D, Tracey I. Multiple Somatotopic Representations of Heat and Mechanical Pain in the Operculo-Insular Cortex: A High-Resolution fMRI Study. Journal of Neurophysiology. 2010;104: 2863– 2872. doi:10.1152/jn.00253.2010

50. Brooks JCW, Zambreanu L, Godinez A, Craig AD (Bud), Tracey I. Somatotopic organisation of the human insula to painful heat studied with high resolution functional imaging. NeuroImage. 2005;27: 201–209. doi:10.1016/j.neuroimage.2005.03.041

51. Henderson LA, Gandevia SC, Macefield VG. Somatotopic organization of the processing of muscle and cutaneous pain in the left and right insula cortex: A single-trial fMRI study. Pain. 2007;128: 20–30. doi:10.1016/j.pain.2006.08.013

52. Gil-Lievana E, Balderas I, Moreno-Castilla P, Luis-Islas J, McDevitt RA, Tecuapetla F, et al. Glutamatergic basolateral amygdala to anterior insular cortex circuitry maintains rewarding contextual memory. Commun Biol. 2020;3: 139. doi:10.1038/s42003-020-0862-z

53. Ghaziri J, Tucholka A, Girard G, Boucher O, Houde J-C, Descoteaux M, et al. Subcortical structural connectivity of insular subregions. Sci Rep. 2018;8: 8596. doi:10.1038/s41598-018-26995-0

54. Bastuji H, Frot M, Perchet C, Hagiwara K, Garcia-Larrea L. Convergence of sensory and limbic noxious input into the anterior insula and the emergence of pain from nociception. Sci Rep. 2018;8: 13360. doi:10.1038/s41598-018-31781-z

55. Uddin LQ. Salience processing and insular cortical function and dysfunction. Nat Rev Neurosci. 2015;16: 55–61. doi:10.1038/nrn3857

56. Seeley WW. The Salience Network: A Neural System for Perceiving and Responding to Homeostatic Demands. J Neurosci. 2019;39: 9878–9882. doi:10.1523/JNEUROSCI.1138-17.2019

57. Benarroch EE. The Central Autonomic Network: Functional Organization, Dysfunction, and Perspective. Mayo Clinic Proceedings. 1993;68: 988–1001. doi:10.1016/S0025-6196(12)62272-1

58. Quadt L, Critchley H, Nagai Y. Cognition, emotion, and the central autonomic network. Autonomic Neuroscience. 2022;238: 102948. doi:10.1016/j.autneu.2022.102948

59. Craig AD. How do you feel? Interoception: the sense of the physiological condition of the body. Nat Rev Neurosci. 2002;3: 655–666. doi:10.1038/nrn894

60. Khalsa SS, Adolphs R, Cameron OG, Critchley HD, Davenport PW, Feinstein JS, et al. Interoception and Mental Health: A Roadmap. Biological Psychiatry: Cognitive Neuroscience and Neuroimaging. 2018;3: 501–513. doi:10.1016/j.bpsc.2017.12.004

61. Verdejo-Garcia A, Clark L, Dunn BD. The role of interoception in addiction: A critical review. Neuroscience & Biobehavioral Reviews. 2012;36: 1857–1869. doi:10.1016/j.neubiorev.2012.05.007

62. Downar J, Blumberger DM, Daskalakis ZJ. The Neural Crossroads of Psychiatric Illness: An Emerging Target for Brain Stimulation. Trends in Cognitive Sciences. 2016;20: 107–120. doi:10.1016/j.tics.2015.10.007

63. Namkung H, Kim S-H, Sawa A. The Insula: An Underestimated Brain Area in Clinical Neuroscience, Psychiatry, and Neurology. Trends in Neurosciences. 2017;40: 200–207. doi:10.1016/j.tins.2017.02.002

64. Goodkind M, Eickhoff SB, Oathes DJ, Jiang Y, Chang A, Jones-Hagata LB, et al. Identification of a Common Neurobiological Substrate for Mental Illness. JAMA Psychiatry. 2015;72: 305–315. doi:10.1001/jamapsychiatry.2014.2206

65. Palma J-A, Benarroch EE. Neural control of the heart: Recent concepts and clinical correlations. Neurology. 2014;83: 261–271. doi:10.1212/WNL.0000000000000605

66. Oppenheimer S, Cechetto D. The Insular Cortex and the Regulation of Cardiac Function. 1st ed. In: Terjung R, editor. Comprehensive Physiology. 1st ed. Wiley; 2016. pp. 1081–1133. doi:10.1002/cphy.c140076

67. Oppenheimer SM, Gelb A, Girvin JP, Hachinski VC. Cardiovascular effects of human insular cortex stimulation. Neurology. 1992;42: 1727–1727. doi:10.1212/WNL.42.9.1727

68. Chouchou F, Mauguière F, Vallayer O, Catenoix H, Isnard J, Montavont A, et al. How the insula speaks to the heart: Cardiac responses to insular stimulation in humans. Hum Brain Mapp. 2019;40: 2611–2622. doi:10.1002/hbm.24548

69. Liberati G, Mulders D, Algoet M, van den Broeke EN, Santos SF, Ribeiro Vaz JG, et al. Insular responses to transient painful and non-painful thermal and mechanical spinothalamic stimuli recorded using intracerebral EEG. Sci Rep. 2020;10: 22319. doi:10.1038/s41598-020-79371-2

70. Pollatos O, Füstös J, Critchley HD. On the generalised embodiment of pain: How interoceptive sensitivity modulates cutaneous pain perception. PAIN®. 2012;153: 1680– 1686. doi:10.1016/j.pain.2012.04.030

71. Paulus MP, Stewart JL. Interoception and drug addiction. Neuropharmacology. 2014;76: 342–350. doi:10.1016/j.neuropharm.2013.07.002

72. Paulus MP, Stein MB. An Insular View of Anxiety. Biological Psychiatry. 2006;60: 383–387. doi:10.1016/j.biopsych.2006.03.042

73. Bonaz B, Lane RD, Oshinsky ML, Kenny PJ, Sinha R, Mayer EA, et al. Diseases, Disorders, and Comorbidities of Interoception. Trends in Neurosciences. 2021;44: 39–51. doi:10.1016/j.tins.2020.09.009

74. Brewer R, Murphy J, Bird G. Atypical interoception as a common risk factor for psychopathology: A review. Neuroscience & Biobehavioral Reviews. 2021;130: 470–508. doi:10.1016/j.neubiorev.2021.07.036

75. Rossi S, Antal A, Bestmann S, Bikson M, Brewer C, Brockmöller J, et al. Safety and recommendations for TMS use in healthy subjects and patient populations, with updates on training, ethical and regulatory issues: Expert Guidelines. Clinical Neurophysiology. 2021;132: 269–306. doi:10.1016/j.clinph.2020.10.003

76. Beck AT, Steer RA, Brown G. Beck Depression Inventory–II. 1996. doi:10.1037/t00742-000

77. Beck AT, Epstein N, Brown G, Steer R. Beck Anxiety Inventory. 1988. doi:10.1037/t02025-000

78. Spielberger CD. State-Trait Anxiety Inventory for Adults. 1983. doi:10.1037/t06496-000

79. Sullivan MJL, Bishop SR, Pivik J. The Pain Catastrophizing Scale: Development and validation. Psychological Assessment. 1995;7: 524–532. doi:10.1037/1040-3590.7.4.524

80. Kroenke K, Spitzer RL, Williams JBW. The Patient Health Questionnaire-2: validity of a two-item depression screener. Med Care. 2003;41: 1284–1292. doi:10.1097/01.MLR.0000093487.78664.3C

81. Taylor JM. Psychometric analysis of the Ten-Item Perceived Stress Scale. Psychol Assess. 2015;27: 90–101. doi:10.1037/a0038100

82. Legon W, Adams S, Bansal P, Patel PD, Hobbs L, Ai L, et al. A retrospective qualitative report of symptoms and safety from transcranial focused ultrasound for neuromodulation in humans. Sci Rep. 2020;10: 5573. doi:10.1038/s41598-020-62265-8

83. Strohman A, In A, Stebbins K, Legon W. Evaluation of a Novel Acoustic Coupling Medium for Human Low-Intensity Focused Ultrasound Neuromodulation Applications. Ultrasound in Medicine & Biology. 2023 [cited 15 Mar 2023]. doi:10.1016/j.ultrasmedbio.2023.02.003

84. Guo H, Hamilton II M, Offutt SJ, Gloeckner CD, Li T, Kim Y, et al. Ultrasound produces extensive brain activation via a cochlear pathway. Neuron. 2018;98: 1020–1030.

85. Boutet A, Gwun D, Gramer R, Ranjan M, Elias GJB, Tilden D, et al. The relevance of skull density ratio in selecting candidates for transcranial MR-guided focused ultrasound. Journal of Neurosurgery. 2019;132: 1785–1791. doi:10.3171/2019.2.JNS182571

86. D’Souza M, Chen KS, Rosenberg J, Elias WJ, Eisenberg HM, Gwinn R, et al. Impact of skull density ratio on efficacy and safety of magnetic resonance–guided focused ultrasound treatment of essential tremor. Journal of Neurosurgery. 2019;132: 1392–1397. doi:10.3171/2019.2.JNS183517

87. Maris E, Oostenveld R. Nonparametric statistical testing of EEG- and MEG-data. Journal of Neuroscience Methods. 2007;164: 177–190. doi:10.1016/j.jneumeth.2007.03.024

88. Chen ACN, Niddam DM, Arendt-Nielsen L. Contact heat evoked potentials as a valid means to study nociceptive pathways in human subjects. Neuroscience Letters. 2001;316: 79–82. doi:10.1016/S0304-3940(01)02374-6

89. De Schoenmacker I, Berry C, Blouin J-S, Rosner J, Hubli M, Jutzeler CR, et al. An intensity matched comparison of laser- and contact heat evoked potentials. Sci Rep. 2021;11: 6861. doi:10.1038/s41598-021-85819-w

90. Treeby BE, Cox BT. k-Wave: MATLAB toolbox for the simulation and reconstruction of photoacoustic wave fields. JBO. 2010;15: 021314. doi:10.1117/1.3360308

91. Mueller JK, Ai L, Bansal P, Legon W. Computational exploration of wave propagation and heating from transcranial focused ultrasound for neuromodulation. Journal of neural engineering. 2016;13: 056002.

92. Aubry J-F, Tanter M, Pernot M, Thomas J-L, Fink M. Experimental demonstration of noninvasive transskull adaptive focusing based on prior computed tomography scans. The Journal of the Acoustical Society of America. 2003;113: 84–93. doi:10.1121/1.1529663

93. Marquet F, Pernot M, Aubry J-F, Montaldo G, Marsac L, Tanter M, et al. Non-invasive transcranial ultrasound therapy based on a 3D CT scan: protocol validation and$\less$i$\greater$in vitro$\less$/i$\greater$results. Phys Med Biol. 2009;54: 2597– 2613. doi:10.1088/0031-9155/54/9/001

94. Lenhard W, Lenhard A. Testing the Significance of Correlations. 2014 [cited 21 Mar 2023]. doi:10.13140/RG.2.1.2954.1367

95. van Wijk G, Veldhuijzen DS. Perspective on Diffuse Noxious Inhibitory Controls as a Model of Endogenous Pain Modulation in Clinical Pain Syndromes. The Journal of Pain. 2010;11: 408–419. doi:10.1016/j.jpain.2009.10.009

96. Coulombe M-A, Erpelding N, Kucyi A, Davis KD. Intrinsic functional connectivity of periaqueductal gray subregions in humans. Human Brain Mapping. 2016;37: 1514–1530. doi:10.1002/hbm.23117

97. Lee J-Y, You T, Lee C-H, Im GH, Seo H, Woo C-W, et al. Role of anterior cingulate cortex inputs to periaqueductal gray for pain avoidance. Current Biology. 2022;32: 2834–2847.e5. doi:10.1016/j.cub.2022.04.090

98. Wang AL, Mouraux A, Liang M, Iannetti GD. Stimulus Novelty, and Not Neural Refractoriness, Explains the Repetition Suppression of Laser-Evoked Potentials. Journal of Neurophysiology. 2010;104: 2116–2124. doi:10.1152/jn.01088.2009

99. Ronga I, Valentini E, Mouraux A, Iannetti GD. Novelty is not enough: laser-evoked potentials are determined by stimulus saliency, not absolute novelty. Journal of Neurophysiology. 2013;109: 692–701. doi:10.1152/jn.00464.2012

100. Downar J, Crawley AP, Mikulis DJ, Davis KD. A multimodal cortical network for the detection of changes in the sensory environment. Nat Neurosci. 2000;3: 277–283. doi:10.1038/72991

101. Menon V, Uddin LQ. Saliency, switching, attention and control: a network model of insula function. Brain Struct Funct. 2010;214: 655–667. doi:10.1007/s00429-010-0262-0

102. Segerdahl AR, Mezue M, Okell TW, Farrar JT, Tracey I. The dorsal posterior insula is not an island in pain but subserves a fundamental role - Response to: “Evidence against pain specificity in the dorsal posterior insula” by Davis et al. F1000Res. 2015;4: 1207. doi:10.12688/f1000research.7287.1

103. Davis KD, Bushnell MC, Iannetti GD, St. Lawrence K, Coghill R. Evidence against pain specificity in the dorsal posterior insula. F1000Res. 2015;4: 362. doi:10.12688/f1000research.6833.1

104. Bastuji H, Frot M, Perchet C, Hagiwara K, Garcia-Larrea L. Convergence of sensory and limbic noxious input into the anterior insula and the emergence of pain from nociception. Sci Rep. 2018;8: 13360. doi:10.1038/s41598-018-31781-z

105. Lutz A, McFarlin DR, Perlman DM, Salomons TV, Davidson RJ. Altered anterior insula activation during anticipation and experience of painful stimuli in expert meditators. NeuroImage. 2013;64: 538–546. doi:10.1016/j.neuroimage.2012.09.030

106. De Schoenmacker I, Archibald J, Kramer JLK, Hubli M. Improved acquisition of contact heat evoked potentials with increased heating ramp. Sci Rep. 2022;12: 925. doi:10.1038/s41598-022-04867-y

107. Valeriani M, Le Pera D, Niddam D, Chen ACN, Arendt-Nielsen L. Dipolar modelling of the scalp evoked potentials to painful contact heat stimulation of the human skin. Neuroscience Letters. 2002;318: 44–48. doi:10.1016/S0304-3940(01)02466-1

108. Shenoy R, Roberts K, Papadaki A, McRobbie D, Timmers M, Meert T, et al. Functional MRI brain imaging studies using the Contact Heat Evoked Potential Stimulator (CHEPS) in a human volunteer topical capsaicin pain model. JPR. 2011;4: 365–371. doi:10.2147/JPR.S24810

109. Barrett LF, Simmons WK. Interoceptive predictions in the brain. Nat Rev Neurosci. 2015;16: 419–429. doi:10.1038/nrn3950

110. Başar E. EEG — Dynamics and Evoked Potentials in Sensory and Cognitive Processing by the Brain. In: Başar E, editor. Dynamics of Sensory and Cognitive Processing by the Brain. Berlin, Heidelberg: Springer; 1988. pp. 30–55. doi:10.1007/978-3-642-71531-0_3

111. Kolev V, Yordanova J. Analysis of phase-locking is informative for studying event-related EEG activity. Biol Cybern. 1997;76: 229–235. doi:10.1007/s004220050335

112. Cheron G, Cebolla AM, De Saedeleer C, Bengoetxea A, Leurs F, Leroy A, et al. Pure phase-locking of beta/gamma oscillation contributes to the N30 frontal component of somatosensory evoked potentials. BMC Neuroscience. 2007;8: 75. doi:10.1186/1471-2202-8-75

113. Kim JA, Davis KD. Neural Oscillations: Understanding a Neural Code of Pain. Neuroscientist. 2021;27: 544–570. doi:10.1177/1073858420958629

114. Harris HN, Peng YB. Evidence and explanation for the involvement of the nucleus accumbens in pain processing. Neural Regen Res. 2019;15: 597–605. doi:10.4103/1673-5374.266909

115. Knyazev GG. EEG delta oscillations as a correlate of basic homeostatic and motivational processes. Neuroscience & Biobehavioral Reviews. 2012;36: 677–695. doi:10.1016/j.neubiorev.2011.10.002

116. Brandenberger G, Ehrhart J, Piquard F, Simon C. Inverse coupling between ultradian oscillations in delta wave activity and heart rate variability during sleep. Clinical Neurophysiology. 2001;112: 992–996. doi:10.1016/S1388-2457(01)00507-7

117. Ako M, Kawara T, Uchida S, Miyazaki S, Nishihara K, Mukai J, et al. Correlation between electroencephalography and heart rate variability during sleep. Psychiatry and Clinical Neurosciences. 2003;57: 59–65. doi:10.1046/j.1440-1819.2003.01080.x

118. Ploner M, Sorg C, Gross J. Brain Rhythms of Pain. Trends in Cognitive Sciences. 2017;21: 100–110. doi:10.1016/j.tics.2016.12.001

119. Kemp AH, Quintana DS. The relationship between mental and physical health: Insights from the study of heart rate variability. International Journal of Psychophysiology. 2013;89: 288–296. doi:10.1016/j.ijpsycho.2013.06.018

120. Thayer JF, Sternberg E. Beyond Heart Rate Variability. Annals of the New York Academy of Sciences. 2006;1088: 361–372. doi:10.1196/annals.1366.014

121. Duck FA. Medical and non-medical protection standards for ultrasound and infrasound. Progress in Biophysics and Molecular Biology. 2007;93: 176–191. doi:10.1016/j.pbiomolbio.2006.07.008

122. Health C for D and R. Marketing Clearance of Diagnostic Ultrasound Systems and Transducers. In: U.S. Food and Drug Administration [Internet]. FDA; 27 Jun 2019 [cited 25 Jan 2022]. Available: https://www.fda.gov/regulatory-information/search-fda-guidance-documents/marketing-clearance-diagnostic-ultrasound-systems-and-transducers

